# Molecular Transfer Model for pH effects on Intrinsically Disordered Proteins: Theory and Applications

**DOI:** 10.1101/2020.12.02.408849

**Authors:** Mauro L. Mugnai, D. Thirumalai

## Abstract

We present a theoretical method to study how changes in pH shape the heterogeneous conformational ensemble explored by intrinsically disordered proteins (IDPs). The theory is developed in the context of coarse-grained models, which enable a fast, accurate, and extensive exploration of conformational space at a given protonation state. In order to account for pH effects, we generalize the Molecular Transfer Model (MTM), in which conformations are re-weighted using the transfer free energy, which is the free energy necessary for bringing to equilibrium in a new environment a “frozen” conformation of the system. Using the semi-grand ensemble, we derive an exact expression of the transfer free energy, which amounts to the appropriate summation over all the protonation states. Because the exact result is computationally too demanding to be useful for large polyelectrolytes or IDPs, we introduce a mean-field (MF) approximation of the transfer free energy. Using a lattice model, we compare the exact and MF results for the transfer free energy and a variety of observables associated with the model IDP. We find that the precise location of the charged groups (the sequence), and not merely the net charge, determines the structural properties. We demonstrate that some of the limitations previously noted for MF theory in the context of globular proteins are mitigated when disordered polymers are studied. The excellent agreement between the exact and MF results poises us to use the method presented here as a computational tool to study the properties of IDPs and other biological systems as a function of pH.

## Introduction

The cellular environment spans a broad spectrum of solution acidity/basicity. For instance, the vesicles of the secretory and endocytic pathways,^1^ melanosomes, ^2^ and mytochondria^3^ are more acidic than the cytosol, with organelle-dependent differences that are as large as three pH units. In addition, external stimuli could alter the pH, thereby producing dramatic changes in the physiology of the cell. ^4^ At a molecular level, changes in pH alter the structure and solubility of proteins, and promote the formation of condensates,^5–7^ and fibrils,,^8,9^ and could be used tune between these two scenarios.^10,11^ These effects are caused by changes in electrostatic interactions.

Titratable groups (the N- and C-termini, and the side-chains of lysine, arginine, histidine, aspartic acid, glutamic acid, tyrosine, and cysteine) respond to increase (decrease) of pH by releasing (binding) protons. Changes in the charge of the individual groups and the whole protein modulate the strength of both intra-molecular and inter-molecular interactions. In order to understand these effects, it is crucial to estimate the pH “scale” around which the protonation and deprotonation occur, that is the pKa-s. These properties depend (i) on the chemistry of the individual titratable groups, and (ii) on the local environment as well as the protein conformation. We focus here on the strategies to account for the effect of the interaction of all the ionizable groups on the protonation of each one of them, assuming that the properties of titratable groups in isolation are accurately measured.

Over the years, a number of methods have been introduced to account for the role of protein structure and flexibility upon titration. These range from the theoretical model of Tanford and Kirkwood (TK),^12^ to a variety of computational strategies that employ all-atom or coarse-grained description of protein and the solvent. The challenge in devising an accurate strategy to investigate pH effect is clear: for each titratable group there are two possible states, protonated and deprotonated. In sufficiently large proteins, there could be dozens of such amino acids, making it difficult to account exhaustively for all the possible protonation states. Because interactions between the ionized groups affect the titration, whenever the system of interest explores a heterogeneous set of structures, the interplay between conformation and protonation exacerbates the challenge. This is the case for pH-induced unfolding of globular proteins, or for predicting the conformations of IDPs with a large number of residues.^13^ In these cases, successfully accounting for all the relevant protonation states for each sampled structure at a given pH becomes computationally prohibitive.

In computational models with all-atom resolution of the protein, there are two strategies that are used to perform constant pH (as opposed to constant protonation state) simulations: the protonation state is described either by a discrete or by a continuous variable.^14–27^ In the first case, an attempt at adding or removing protons from titratable groups is accomplished by introducing Monte Carlo steps in the protonation space; in the second case the “degree of protonation” of a base or an acid follows a specific time evolution, which can be biased in order to favor fully ionized or neutral states as a means to reduce the population of intermediate “degrees of protonation”, deemed to be unphysical. These strategies yield a simultaneous exploration of conformational and protonation space, which enables the calculation of average properties while retaining dynamic information. However, the calculations must be repeated for each pH value.

An alternative approach, based on the Molecular Transfer Model (MTM),^28,29^ was proposed by O’Brien, Brooks, and Thirumalai (OBT).^30^ The basis of the theory is rooted in the studies of Tanford who estimated the differences in the stability of folded proteins upon a change in solution conditions. Tanford defined the transfer free energy as the free energy cost of transferring the residues and the peptide backbone from one solution condition to another. The free energy for the whole protein could be taken to be the sum of the changes in the transfer free energies of the individual amino acids as well as the peptide backbone upon a change in the solution conditions. Despite the use of the additivity assumption, the Tanford method works remarkably well in obtaining the stability changes of folded proteins as the denaturant concentration is changed. In the OBT approach, the protein is described at a coarse-grained (CG) level in order to extensively sample its conformations, and a model analogous to Tanford’s was introduced to calculate the free energy change when each protein conformation is transferred to a different pH. The OBT theory accounts for the changes in the solvation of titratable groups upon folding/unfolding, which results in sizable modification of the pKa-s of ionizable residues depending on the overall conformation of the protein. An advantage of this model is that once an extensive simulation is conducted at a given pH, features computed in different solution conditions emerge after a fast re-weighting of the sampled conformations. Despite the success of MTM in quantitatively reproducing experimental results for pH-dependent folding free energy of globular proteins,^28^ subsequent CG strategies resorted to implementing pH effects in the mold of all-atom models, with a “dynamical” instead of a “statistical” approach to sampling protonation states.^31–34^

Here, we reprise the MTM approach of OBT,^30^ and modify it so that it could be used to investigate pH effects on IDPs, which is the eventual goal of this program. In IDPs, the presence of a heterogeneous ensemble of globular and extended states demands a finer sensitivity to the local, transient conformation around each titratable group. To overcome this problem while retaining the simplicity of the MTM, we developed a new method by building on the works of TK,^12^ Tanford and Roxby (TR),^35^ Bashford and Karplus (BK),^14^ Gilson,^15^ and Baptista *et al*.^18^ In a nutshell, we assume that the changes in pKa of each group emerge from a combination of given chemical properties and electrostatic interactions with other charged groups, and neglect the effect of other interactions such as hydrogen bonds, as is commonly done (see for instance the work of TK^12^). With this assumption, we use the semi-grand ensemble^18,36^ to derive a transfer free energy. In this way, we recover an expression that was previously reported by BK^14^ and Gilson^15^ on the basis of thermodynamic considerations, starting from either a fully de-protonated or fully uncharged protein (see the Supporting Information (SI)). The exact calculation of the pH-dependent transfer free energy is not feasible for systems with more than a dozen titratable groups. Therefore, it is tempting to adopt a mean-field (MF) approximation.^14^ However, BK^14^ and Gilson^15^ argued that the MF approximation is inaccurate when there are strong interactions between nearby titratable groups. Therefore, they attempted to account for the large number of protonation states by introducing alternative strategies, such as the reduced-site method, in which only a subset of titratable groups can be protonated/deprotonated at a given pH,^14^ or a technique based on clustering in order to separate local interactions (treated exactly) from long-range interactions (approximated using MF). ^15^ Notably, in these influential works, BK and Gilson did not explore the effect of different conformations to address the accuracy of the MF approximation. The protein conformation was quenched to the native structure. Of course, fixing the conformation in IDPs would produce qualitatively incorrect results.

We hypothesized that the inaccuracies in the MF theory would be less prominent for IDPs, which do not have persistent and stable globular structures. We expect titratable groups of the IDPs to be well-solvated, and consequently interactions between the charged residues are weakened by the ionic strength and high dielectric permittivity of the solution. To test our conjecture, which has similarities with ideas proposed in the context of the unfolded state,^17^ we constructed a simple and precisely solvable cubic lattice model of an IDP. The simplicity of the model permits exhaustive enumeration of all the conformations and protonation states, thereby allowing us to test the validity of the MF approximation without worrying about sampling issues. We show that the agreement between the numerically exact results and those obtained using the MF approximation is excellent. Finally, we show that the transfer free energy of OBT^30^ used to investigate pH effects on folding of globular proteins is recovered with the aid of a simple argument (see the SI). Our findings pave the way to use the MTM for IDPs (MTM-IDP) to probe pH effects in a variety of problems including, but not limited to, the IDPs.

## Methods

### Anions, Cations, Protonation, and Notation

We first introduce some notation in order to make the rest of the Methods section easier to understand. As we discuss the nomenclature, we focus on proteins, especially IDPs. However, the general scheme holds for RNA, DNA, polyelectrolytes, polyampholytes as well as small molecules.

Let a protein have *N*_*T*_ titratable groups, which are capable of binding or releasing protons according to the following reaction,

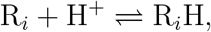

where R_*i*_ is the *i*-th titratable chemical species and H^+^ refers to the proton. We divide these species into two classes. We refer to the N-terminal amino group, and the side-chains of histidine, lysine, and arginine as “cataionic groups” because in their ionized form they are positively charged. In contrast, “anionic groups” such as the carboxyl C-terminal end, and the side-chains of the aspartic acid, glutamic acid, tyrosine, and cysteine are negatively charged when ionized. Cationic groups become positively charged in the protonated state; in contrast, anionic groups are neutral when they bind a proton.

In order to distinguish between the protonated (R_*i*_H) and de-protonated (R_*i*_) forms of the titratable group, we introduce the following set of labels: (i) the ionization state *x*_*i*_, which is = 1 if the *i*-th group bears a charge and = 0 otherwise. (ii) The protonation state, *p*_*i*_, is = 1 (= 0) when the group is protonated (de-protonated). (iii) The charge of a group, *q*_*i*_, which is equal to *z*_*i*_ *· x*_*i*_, where *z*_*i*_ = 1 for cationic groups, and *z*_*i*_ = −1 for anionic groups. The values assumed by the labels in the different scenarios are illustrated in Fig. 1. The relationship between protonation states and ionization state is given by,

**Figure 1:**
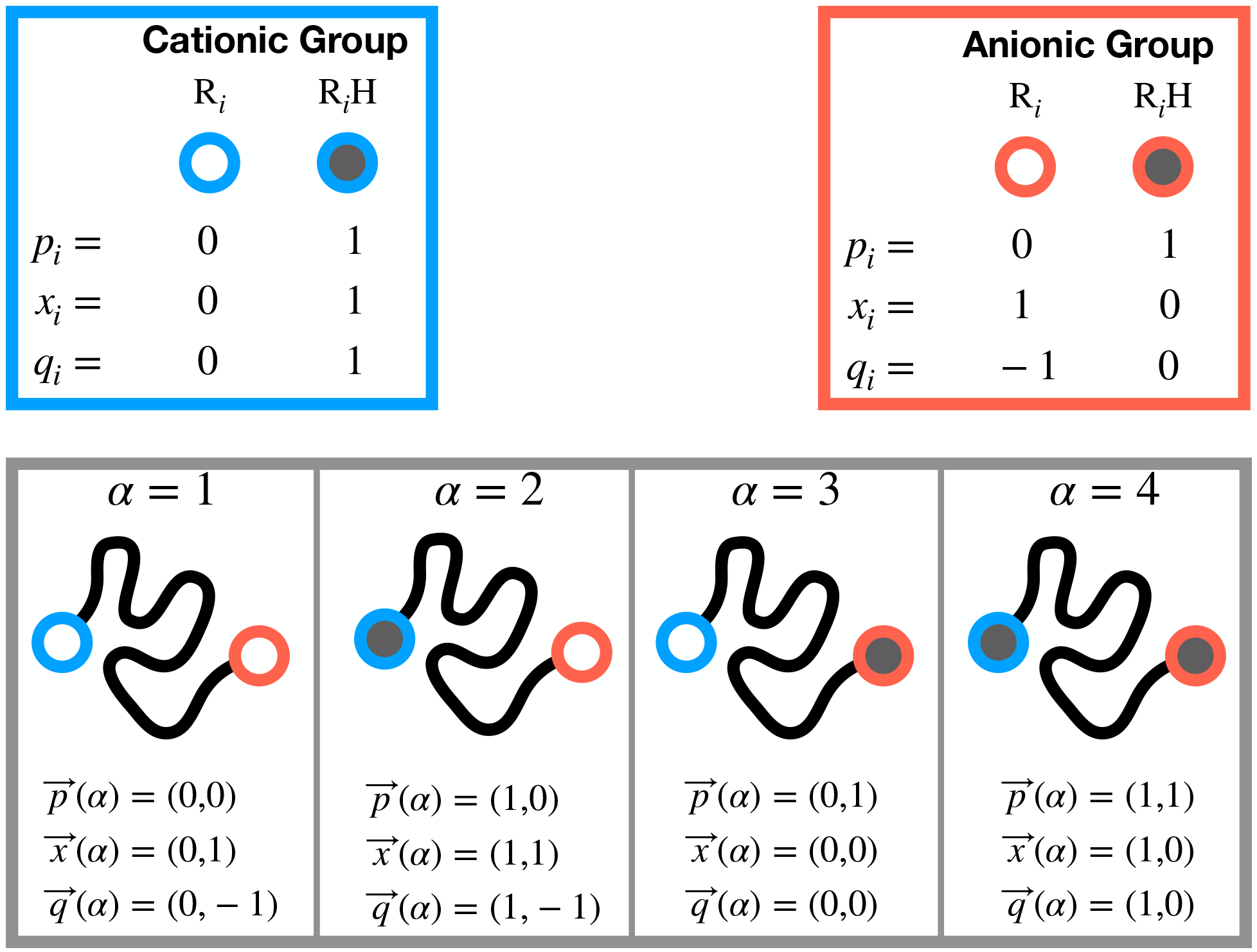
Pictorial representation of ionization and protonation. The top two boxes show the protonation state *p*_*i*_, ionization state *x*_*i*_, and charge *q*_*i*_ for cationic (blue) and anionic (red) groups. Empty dots represent de-protonated states, the gray-filled dots indicates that the group is protonated. The bottom gray box shows all the protonation/ionization states for a frozen conformation of a polymer (black) with two titratable groups: a cationic and an anionic one. Each of the four conformations is labeled with a value of *α* that indicates the distinct protonation/ionization states (4 in this example), and the corresponding protonation 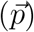 and ionization 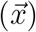 states are shown, together with the charge 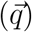.

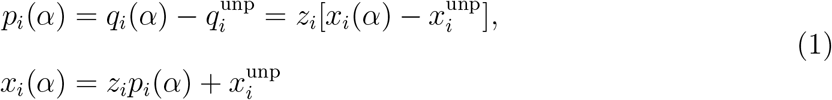

where 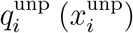 is the charge (ionization state) of the *i*-th group in the un-protonated state. For an anionic group, the charge in the un-protonated state is −1 and the ionization state value is 1, whereas these values are 0 for a cationic group (see Fig. 1).

A protein with *N*_*T*_ titratable groups (*N*_*T*_ = 2 in the example of Fig. 1) has 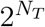 ionization or protonation states (in Fig. 1, 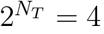), each of which is identified by vectors 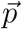 and 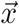 of 0-s and 1-s. We introduce an index *α* which goes from *α* = 1 to 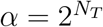. This index corresponds to a unique protonation or ionization state, so that, for instance, 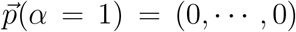 and 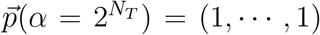, with the corresponding ionization states obtained using the transformation in Eq. 1. The total number of titratable states that are protonated is given by,

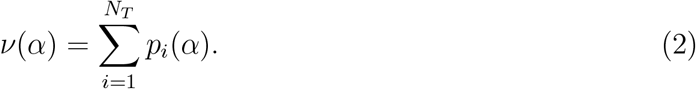

At equilibrium, the dissociation constant in molar (M) units, K_a*i*_, for protonating a R_*i*_ is,

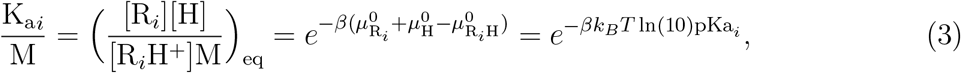

where the symbol [X] refers to the concentration of species X in molar units, and the standard free energies of R_*i*_H, R_*i*_, and the proton are given by 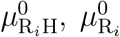 and 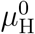 respectively. In Eq. 3 we have also used the definition of pKa of the *i*-th titratable group, which is given by, pKa_*i*_ = − log_10_ K_a*i*_*/*M. The value of this pKa is an intrinsic measurable property of a chemical species measured in isolation and in solution. We assume these values to be given.^37^

### Exact Formulation

We present a general method to compute the transfer free energy, following Tanford’s studies in estimating stability of globular proteins as a function of denaturant concentration. In order to complete this task, it proves convenient to use the semi-grand canonical ensemble.^36^ The partition function in this ensemble is:

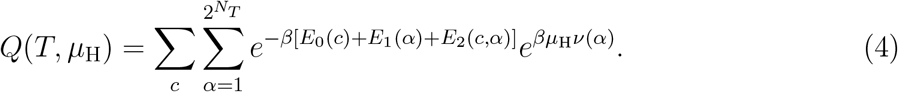

In the above equation, *c* labels the spatial conformation (it could be replaced by a continuous index, but we keep it discrete without loss of generality), *E*_0_(*c*) is an energy term which depends only on the conformation, *E*_1_(*α*) is determined solely by the protonation of titratable groups, and *E*_2_(*c, α*) couples protonation and spatial conformation. Note that we did not include in the partition function the interaction between free protons, which is tantamount to assuming that their behavior is ideal. Thus, we can approximate the proton chemical potential, *µ*_H_, in Eq. 4 as,

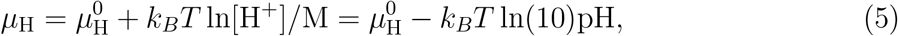

where pH = − log_10_[H^+^]*/*M. Note that we have excluded solvent and salt effects from the partition function; in the coarse grained representation of the polymer their contribution is to modulate the effective energies, *E*_0_(*c*), *E*_1_(*α*), and *E*_2_(*c, α*), which will become clear later. There is no pressure (or volume) term in the partition function in Eq. 4; its contribution would appear through the energy function and the standard chemical potential of the proton. Under the reasonable assumption that the fluid is incompressible, the results would not be affected by the choice of the thermodynamic variable. In any event, these are standard assumptions in the CG models of proteins^38^ and IDPs.^13,39,40^ They could be relaxed at the cost of increasing the complexity of the calculations without being able to assess the accuracy.

We introduce a reference protonation state, *α*^⋆^, and recast the partition function in the following form,

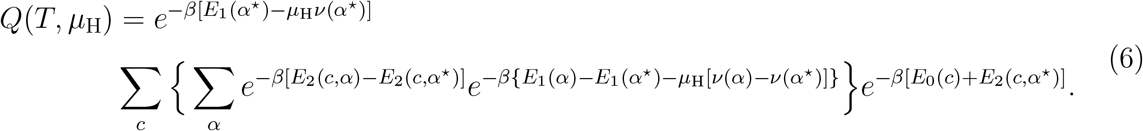

We define the transfer free energy to be,

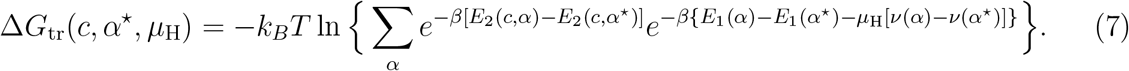

The above equation is evaluated by choosing a specific conformation and summing over *α*, which labels the protonation state of the polymer. Thus, Δ*G*_tr_(*c, α*^⋆^, *µ*_H_) depends only on *c*, pH, and the reference state, *α*^⋆^. Using Eq. 7, the partition function becomes,

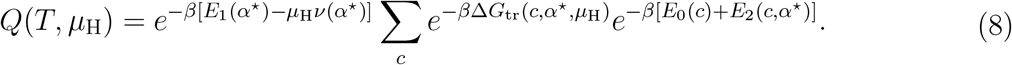

The above expression for *Q*(*T, µ*_H_) has a simple operational interpretation: we perform simulations at a fixed protonation state, *α*^⋆^, and the resulting trajectory provides an ensemble of conformations. To appropriately account for the variations in the protonation state, we re-weight each sampled conformation (fixed *c*) with the corresponding transfer free energy given in Eq. 7. The constant outside the summation in Eq. 8 refers to the titratable groups in state *α*^⋆^. Being a constant, it does not affect the evaluation of average properties of the system, although it does contribute to the absolute value of the total free energy.

*E*_1_(*α*): The transfer free energy in Eq. 7 can be calculated if *E*_1_(*α*) and *E*_2_(*c, α*) are specified. We model *E*_1_(*α*) as follows. Because it is by construction independent on the conformation of the polymer, we assume that it can be written as a sum of individual contributions from each titratable group. Each contribution is modeled as the standard free energy of the protonated or de-protonated form of the *i*-th group isolated in solution,

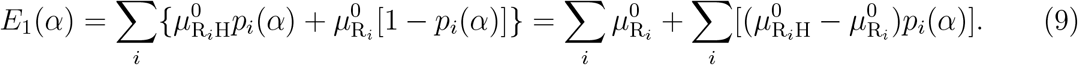

Therefore, recalling the definition of *ν*(*α*) in Eq. 2, the pKa of the *i*-th group in Eq. 3, and the proton chemical potential in Eq. 5, we get,

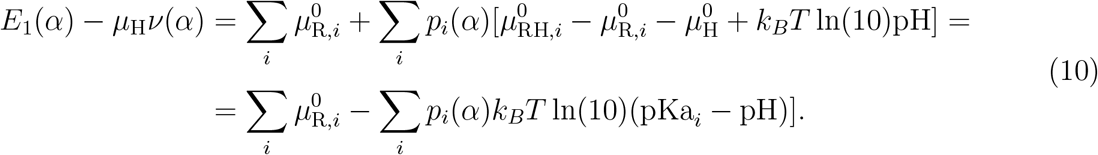

Following the nomenclature of BK^14^ and Gilson,^15^ we introduce *b*_*i*_, defined as,

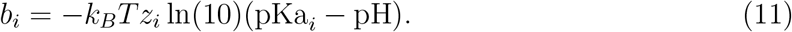

Finally, using Eq. 1 to switch from protonation [*p*_*i*_(*α*)] to ionization [*x*_*i*_(*α*)] states, we write:

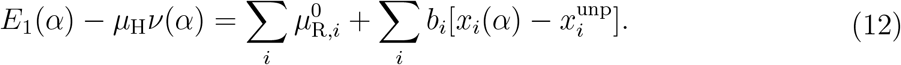

*E*_2_(*c, α*): In order to model to coupling between conformations and protonation states, we assume that the only contribution to *E*_2_(*c, α*) comes from pairwise, electrostatic interactions,^12^ which we write as,

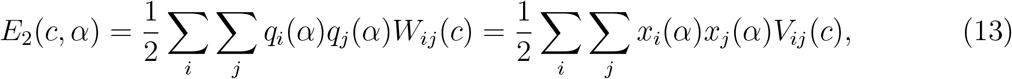

where *z*_*i*_*z*_*j*_*W*_*ij*_(*c*) = *V*_*ij*_(*c*) is an electrostatic interaction depending on the distance between two beads – the functional form will be discussed later. By combining all the terms in the exponent for the transfer free energy (Eq. 7) we get,

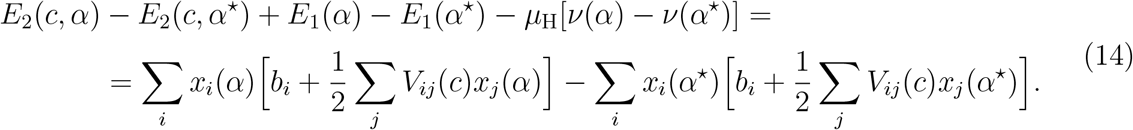

The transfer free energy becomes,

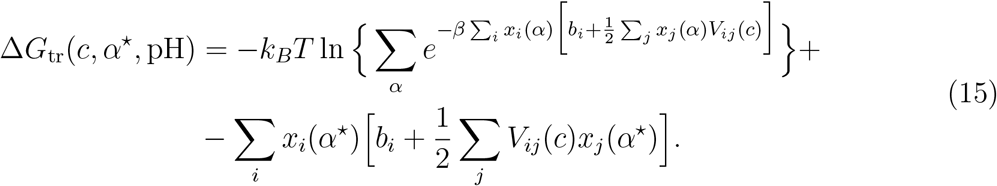

(Note that we changed the argument of the transfer free energy from the chemical potential to the pH). BK^14^ and Gilson^15^ derived results similar to Eq. 15 using two special reference states: the fully de-protonated, and the fully de-ionized (no charges) states, respectively. In the Supporting Information, we show that we can recover the previous results by selecting 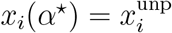and *x*_*i*_(*α*^⋆^) = 0, respectively.

### Mean-field Approximation

The transfer free energy (Eq. 15) is exact for the coarse-grained model provided the effective energies can be decomposed as in Eq. 6, and that they can be accurately accounted for by the expressions that we have chosen for *E*_1_(*α*) and *E*_2_(*c, α*). At this juncture, we need not specify the form of *E*_0_(*c*), which is required only if the method is applied to a specific problem. The calculation of transfer free energy still requires that we explicitly account for all the possible protonation states – a daunting computational task for a protein with a large number of titratable groups.

The computational challenge can be overcome using a MF approximation^14^ to estimate the value of

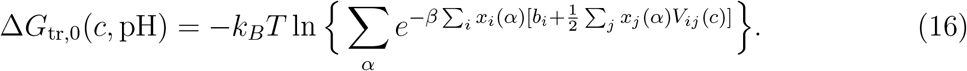

The details of the calculation are the same as those discussed by BK.^14^ The only difference is that BK use protonation instead of ionization states. As a consequence, the final result is the same as the one already provided by Gilson,^15^

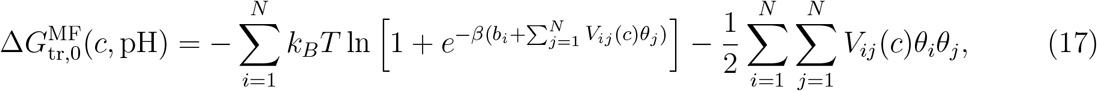

where *θ*_*i*_, the average ionization state of group *i* at a given pH and at a given conformation, and it is equal to,

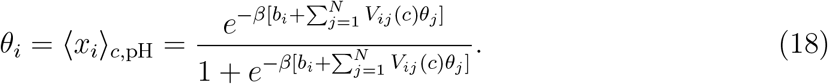

The above equation can be solved using a self-consistent iterative process.^14^ Finally, the MF approximation for the transfer free energy is,

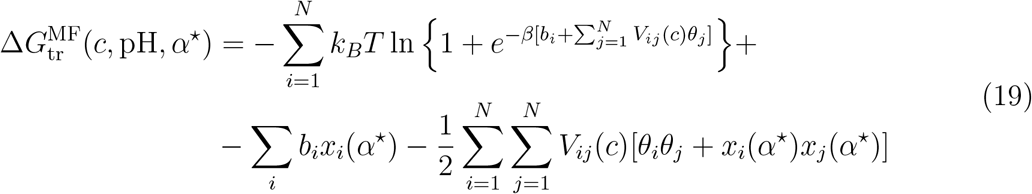

It is easy to show that this result is the same as Gilson’s if the fully de-ionized state [*x*_*i*_(*α*^⋆^) = 0 for all titratable groups]^15^ is chosen as the reference state. Similarly, it is possible to recover the result of BK^14^ if the fully de-protonated 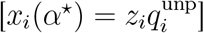 state as the reference.

### Fixed-charge Approximation

An alternative to the MF approximation is to use fixed charges (FC) on each titratable group, which are given by 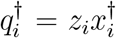, where 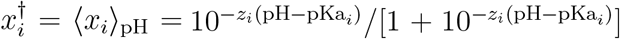. These would be the average charges on each group in isolation, that is in the absence of the other ionizable groups. Note that 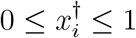, whereas *x*_*i*_ = 0 or = 1. The transfer free energy in this case is given by,

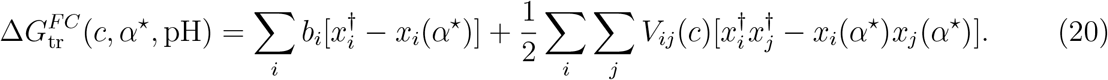

The expression given above is obtained from Eq. 15 by assuming that there is only one ionization state given by, 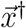.

### Lattice Simulations

In enunciating the principles of protein folding, lattice models with small polymer length for which all allowed conformations could be exactly enumerated, played a vital role.^41^ Here, we tested the accuracy of the MF approximation using a polymer model of length 10 on a simple cubic lattice with spacing, *a* = 3.8 Å. We consider a self-avoiding polymer, for which *E*_0_(*c*) is given by a combination of bonded interactions ensuring that consecutive beads in the sequence occupy nearest-neighbors vertices in the lattice, and excluded volume interactions which forbid two beads from occupying the same lattice site. The functional form of *E*_0_(*c*) is assumed to be such that the energy is zero if these constraints are satisfied (consecutive beads on nearby sites, and the chain does not cross itself), and infinity if they are violated. The total number of allowed conformations is *N*_10mer_ = 1, 853, 886.^42^ To count the conformations we fixed the first bead, and considered conformations related by rotation and specular reflections to be different – more on this later.

The energy *E*_2_(*c, α*) is is modeled as the sum of pairwise Debye-Hückle (DH) electrostatic interactions between charged groups,^36^ where *V*_*ij*_(*c*) is,

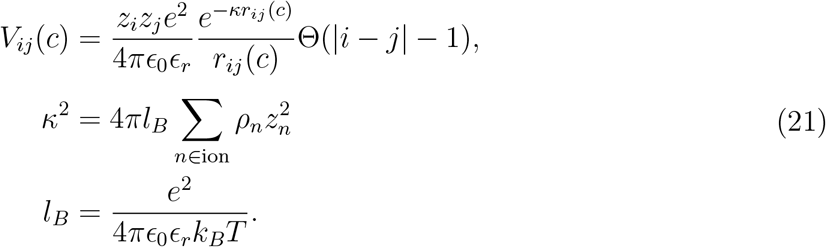

Here, *e* is the unit charge, *E*_0_ is the dielectric vacuum permittivity, *E*_*r*_ is the relative permittivity of the solvent (assumed to be = 78, which is appropriate for an expanded IDP in water). The step function Θ(*x*) (= 1 if *x* > 0 and = 0 otherwise) ensures that only beads that are not consecutive in the chain engage in electrostatic interactions. The inverse of the Debye screening length (*κ*) depends on the Bjerrum length (*l*_*B*_) and on the concentration of screening ions (*ρ*_*n*_), with *z*_*n*_ being the unitless charge on the *n*-th species ion. Finally, 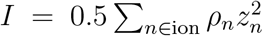 is the ionic strength of the solution. We assume that the solution contains only monovalent ions, so that,

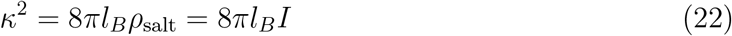

where *ρ*_salt_ = *I* is the sum of the concentrations of dissociated salts.

The pH itself is a measure of the concentration of ions in solution, and must be accounted for when the Debye length is computed. In practice, this means that the sum of the concentration of dissolved salts (monovalent ions), *ρ*_salt_, has two contributions: at low pH (< 7) *ρ*_salt_ = *ρ*_salt,0_ + 10^−pH^M, whereas for high pH (> 7) *ρ*_salt_ = *ρ*_salt,0_ + 10^pH−14^M. Here, 10^−pH^M is the concentration of hydronium ion and a neutralizing (monovalent) anion, 10^pH−14^M is the concentration of hydroxide ion and a neutralizing (monovalent) cation, and *ρ*_salt,0_ is the concentration of any other salt (e.g. NaCl or KCl) such that the ionic strength

is set at the value used for sampling. As a result, given a salt concentration in the DH potential, *ρ*_salt_, the minimum and maximum values of the pH consistent with the simulations are pH = − log_10_(*ρ*_salt_*/*M) and pH = 14 + log_10_(*ρ*_salt_*/*M), respectively, where *ρ*_salt,0_ = 0 M.

### Ensemble Averages

Because the energy function of the polymer on the lattice is invariant upon rotations and reflections, we can combine conformations depending on their geometrical properties. There is one 1-D conformation with degeneracy 6; there are 2033 arrangements of the polymer restricted on a plane with degeneracy 24, and the remaining 3D conformations are 37606 with degeneracy 48, for a total of 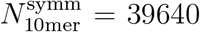 distinguishable conformations (unrelated by symmetry operations), each of which with a weight (*w*) given by the appropriate degeneracy. Using these considerations, the constant-pH average of any observable, *Ô*(*c*), depending solely on the polymer conformation is,

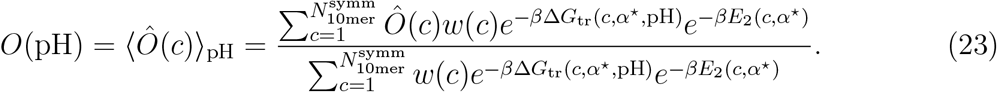

The exact, MF, or FC evaluation of the average of the observable depends on whether we use the exact (Eq. 15), MF (Eq. 19) or FC (Eq. 20) expression for the transfer free energy.

The average of an observable, *Ô*(*α*), which depends on the ionization state alone (for instance the charge on a titratable group) can be written in the following way,

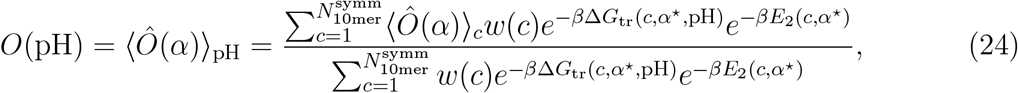

where ⟨*Ô*(*α*)⟩_*c*_ is the average over the ionization computed for a given conformation, that is,

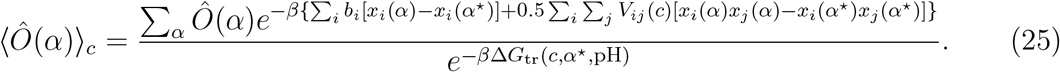

If *Ô*(*α*) is the charge on the *i*-th titratable group, the MF approximation of Eq. 25 is *z*_*i*_*θ*_*i*_, where *θ*_*i*_ is given in Eq. 18. The correlation between charges is given by,

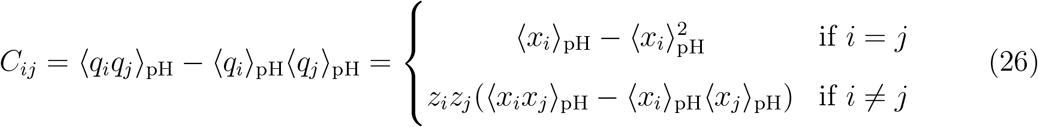

where we used the identities 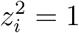 and 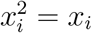.

### Hellinger Distance

In the results section we compare distributions obtained with different approximations for the transfer free energy. In order to quantify the difference between two distributions *q* and *p*, we use the so-called Hellinger distance *H*(*p, q*), given by,

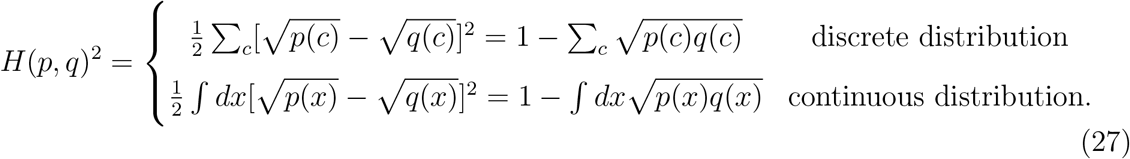

It can be shown that 0 ≤ *H*(*p, q*) *≤* 1, where *H*(*p, q*) = 0 for identical distributions, and *H*(*p, q*) = 1 if the distributions have no overlap.

## Results

We tested the MF (and FC) scheme for the model IDP with the sequence DADAADAAAD, where D stands for aspartic acid (pKa = 4.0^37^), and A is alanine. The purpose is to assess the accuracy of the MF theory. We chose this sequence because it has a high, but not unrealistic density of charged beads. We also distributed the acidic residues along the sequence in such a way that each one of them has a different environment, that is each one of them is followed or preceded by a different number of A-s. Fig. **??** reports the spectrum of the lowest 20 energy levels and the associated degeneracies, as well as the structure of the ground state, and a few of the excited states. The ground state for the fully ionized polymer at all salt concentrations is fully extended. Note that the lowest 20 energy levels are within a *k*_*B*_*T* of each other, indicating that at room temperature they are all populated with nearly identical probability.

### Transfer Free Energy

We computed the exact and MF transfer free energies for all the conformations over a pH range from 2 to 8, and an ionic strength from 1 mM to 1 M (see Fig. 2 and Fig. **??**). As expected, the difference between the transfer free energies is reduced as the ionic strength increases: strong correlations between charged groups make the MF approximation less accurate. As a corollary, increasing the screening is expected to weaken the correlation between charged groups, thus making the MF approximation more accurate. The lowest ionic strength yields the worst results, and presents the only instance in which the MF transfer free energy is smaller (larger in absolute value) than the exact value. Even in this case the differences are less than about 15% (Fig. 2a). At 100mM, which is closer to the physiological concentration, the deviation is no more that 5% at all pH values (Fig. 2b). We also noticed that at low ionic strengths the recursive algorithm necessary to compute the average charges (Eq. 18) does not converge for certain conformations, and leads to an oscillatory solution. We picked the *θ* values resulting in a smaller Δ*G*_tr_ – the rationale is that the MF approximation results from a minimization of the free energy functional in Eq. 16.

**Figure 2:**
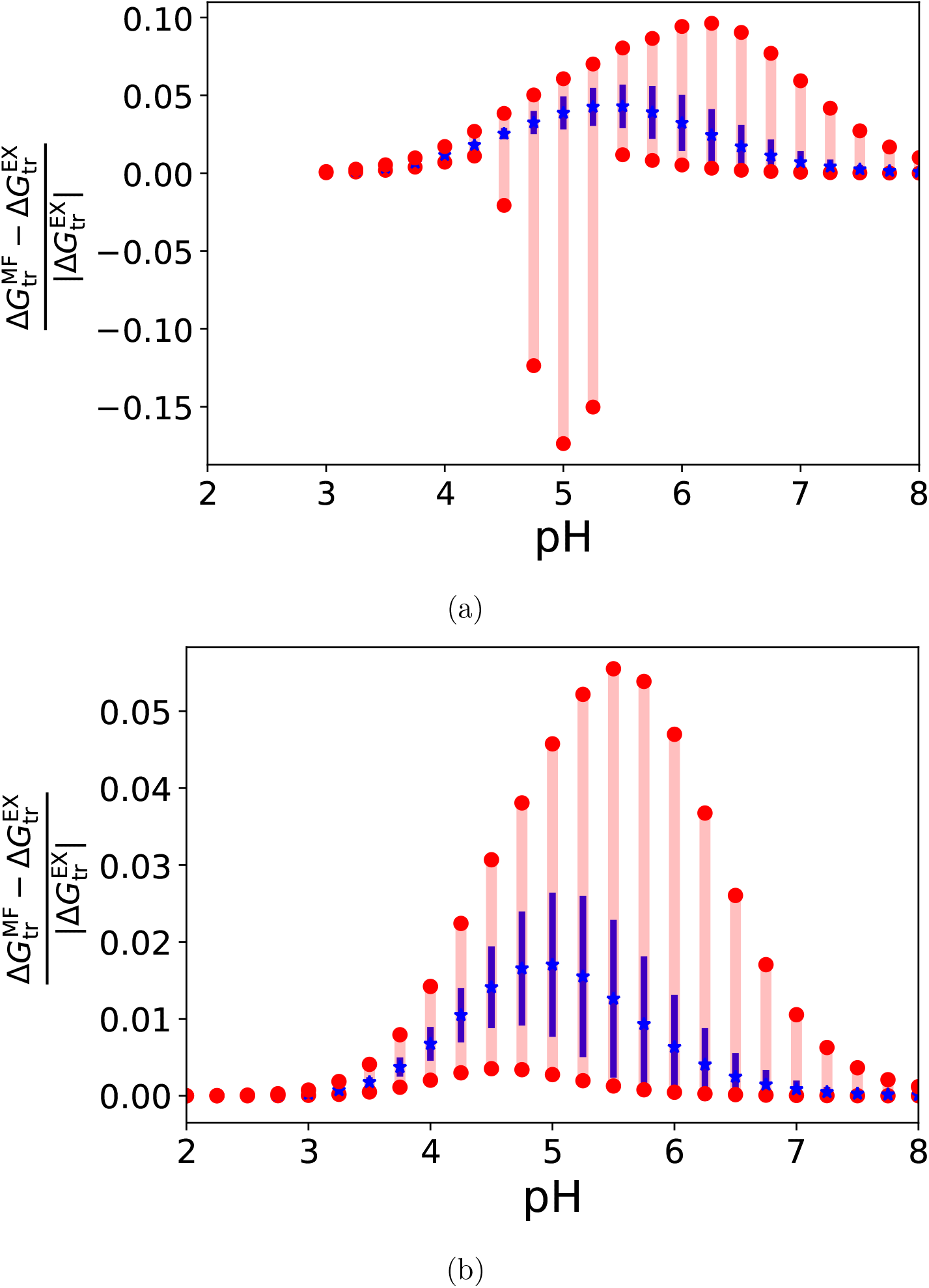
Comparison of the MF and exact calculations of the transfer free energy. The figures show 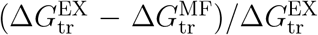 as a function of pH for all sampled conformations unrelated by symmetry operations. The red lines indicate the full range of the distribution, the blue star is the average and the blue bars refer to the standard deviation. The two panels indicate different ionic strengths: (a) 1 mM, (b) 100 mM.

### Probability of Each Conformation

The probability *P* (*k*) a conformation *k*, the mean of any observable *Ô*(*c*) = *δ*_*c,k*_ where *δ* is the Kronecker delta, is calculated using Eq. 23. We computed *P* (*k*) using the exact, MF, and FC ionization states. In Fig. **??** we use the Hellinger distance for discrete distributions (Eq. 27) to show that the MF distribution reproduces the exact result more accurately than the FC. The accuracy increases if pH > pKa or pH < pKa; when pH ≈ pKa there is a peak. The Hellinger distance decreases with the ionic strength of the solution for both MF and FC, indicating that the MF and FC approximations become more reliable as the screening of the electrostatic interaction between the titratable groups increases.

### Average Radius of Gyration

The radius of gyration for a given conformation, *c*, is 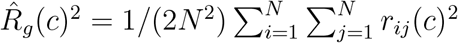, where *r*_*ij*_(*c*) is the distance between two of the *N* (= 10) beads in *c*. The average radius of gyration (*R*_*g*_) is obtained by taking the square root of Eq. 23, where 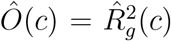 As shown in Fig. 3a, the radii of gyration obtained with the exact and MF evaluations of the transfer free energy are essentially identical. The relative error is less than 1% of the exact value. As the salt concentration increases, the relative discrepancy diminishes, making the MF approximation even more accurate. At strongly acidic pH values, the results are insensitive to salt concentrations: the titratable groups are neutral and the 10-mer is essentially a self-avoiding walk (SAW). The size of the fully charged polymer (high pH, see below) depends on the ionic strength of the solution: as the screening between charges increases, the swelling is less prominent due to the weakening of the repulsion between the ionized groups. Interestingly, the collapse of the polymer, driven by acidification of the solution, is sharper at high ionic strengths: repulsive interactions tend to make the transition less cooperative, likely because they oppose the titration of neighboring anions, which is a correlation effect. At those pH values corresponding to the swelling of the polymer, the FC approximation fails: it underestimates the mid-point of the transition, and overestimates the steepness. The error is somewhat reduced only for the largest ionic strength considered.

**Figure 3:**
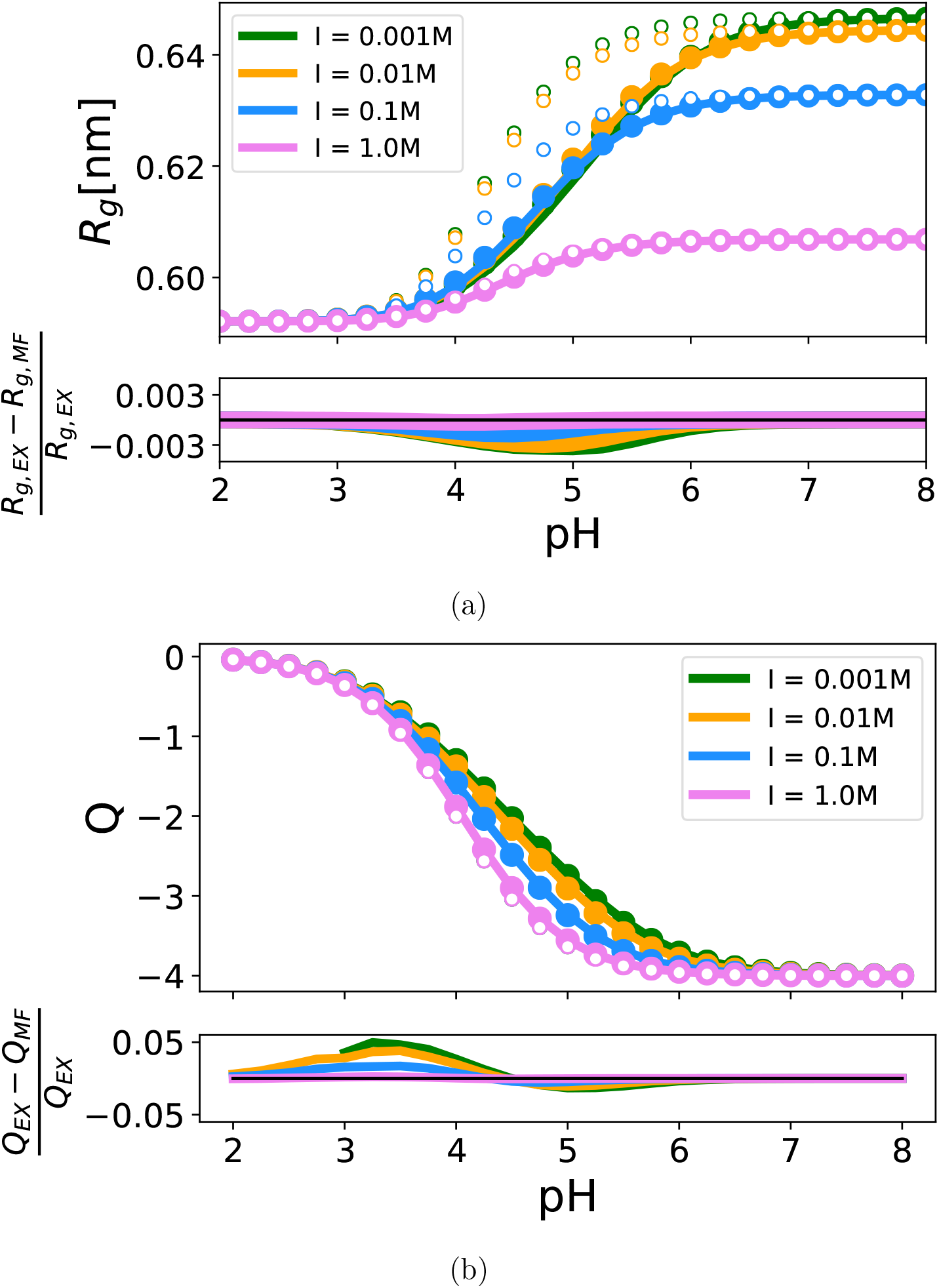
Comparison of the MF, FC, and exact calculations of average titration and radius of gyration. The figure shows *R_g_* (a) and total charge (b) as a function of pH. In the upper panels, lines refer to the exact evaluation of the transfer free energy, dots indicate the results from the MF approximation, and empty dots show the FC results. The bottom panels show the changes of the observables in the exact and MF calculation relative to the exact results. Colors indicate different ionic strengths: green is 1 mM, orange 10 mM, blue 100 mM, and pink 1 M.

### Average Charge

The average charge on each titratable group is given by ⟨*q*_*i*_⟩_pH_ = ⟨*z*_*i*_*x*_*i*_(*α*) ⟩_pH_, where the average is calculated using Eq. 24. The total charge is the sum of the average charge of each group. The average total charge as a function of pH and ionic strength is in Fig. 3b. The maximum relative difference between the exact and MF results is ⪅5% of the exact value, the discrepancy is peaked around pH-values at which the transition occurs, and it diminishes as the ionic strength increases. Notably, the mid point of the transition increases as the salt concentration is diminished, owing to the stronger repulsion between charged groups, which results in an anti-cooperative collective titration (see next section). The increase in the sharpness of the transition with salt concentration confirms this observation. The FC results resemble those obtained for the largest ionic strength, and by construction they are independent of salt concentration, thereby failing to reproduce the exact results within an acceptable error.

### Charge Variance

The correlation between titrating groups is given in Eq. 26. MF methods fail to capture fluctuations, and this is reflected in the fact that the off-diagonal correlation *C*_*ij*_ = 0 (see for instance BK^14^). Similarly, the FC approximation yields no correlation between the charged groups – each residue titrates independently. On the other hand, *C*_*ii*_ is not necessarily zero with either approximation.

In Fig. 4 and Fig. **??** we compare the results of the MF and FC approximations to the exact calculations. The MF approximation is excellent in estimating the diagonal terms of the charge correlations: the location and width of the peaks are correctly identified, with an error that is small at low ionic strength and diminishes at high salt concentrations. At low salt concentrations, the shift of the peak depends on the location of the aspartic acid along the sequence, a feature that is lost with the increase of the ionic strength. In contrast, FC calculations become reliable only at very high salt concentrations, with the four aspartic acids titrating almost independently. The off-diagonal terms are all negative: all charged groups are anionic, thus there is an energetic penalty in charging two aspartic acids simultaneously. The size of the peak of *C*_*ij*_ depends on the distance along the sequence between *i* and *j*: the closer they are, the more they interfere, which results in a larger anti-correlation. The effect is reduced at higher salt concentrations.

**Figure 4:**
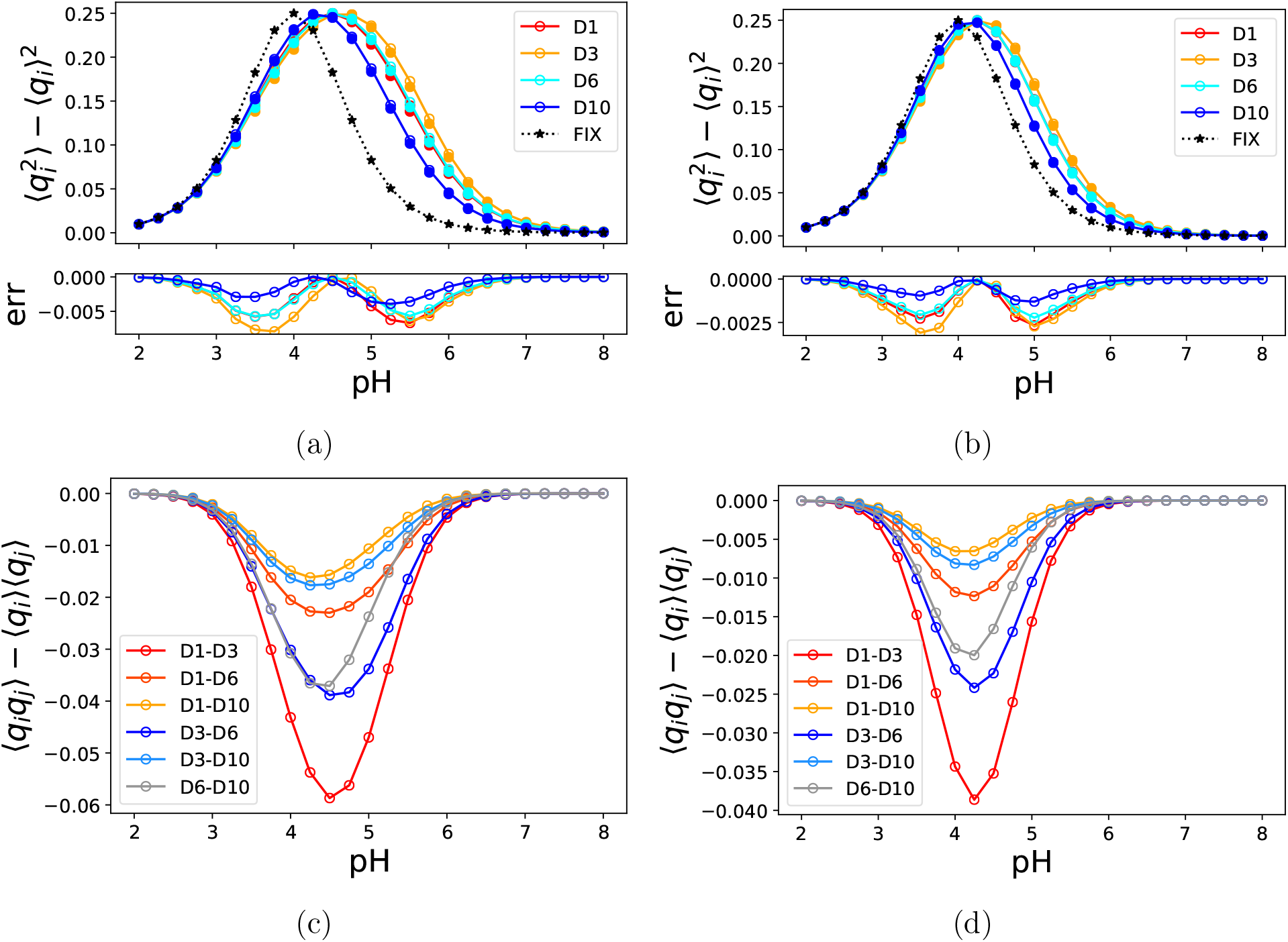
Comparison of the charge-charge correlation as a function of pH. Figures (a) and (b) show the diagonal term of the correlation matrix *C*_*ii*_ in Eq. 26, figures (c) and (d) refer to the off-diagonal terms. Different colors indicate different aspartic acids, as shown in the legends, with empty symbols and lines referring to the exact results, the MF values reported as full circles, and the FC approximation shown as black stars connected by dotted lines. In and (b) the top panels show the correlations, the bottom ones report the error resulting from the MF approximation (MF minus exact). (a) and (c) report the correlations at 1 mM salt concentration, the results at near-physiological ionic strength (100 mM) are shown in panels (b) and (d). Note that in the MF (Hartree-like) approximation *C*_*ij*_ in Eq. 26 is zero if *i* ≠ *j*. The results in (c) and (d) are numerically exact values for the lattice.

### Aspartic Acids are not all Identical

At a given pH, the ionization of each titratable group depends on the conformation, resulting in a distribution of charges. In Fig. 5 and Fig. **??** we show the distribution of charges on the 4 anionic residues as a function of pH and ionic strength. First, we note that although similar, the distributions for the 4 aspartic acids differ: the location along the chain matters. Second, the distributions are sharp at low and high pH values, whereas if the solution pH is in the vicinity of the transition point the distributions become wider. As the solution becomes more acidic, the average of the distribution shifts continuously from −1 to 0. As we observed before, the MF approximation is accurate: the locations and shapes of the exact distributions are captured, and the recovery improves as the salt concentration increases. To make the observations more quantitative, in the insets we report the values of the Hellinger distances, *H*, between the exact and MF distributions (see caption of Fig. 5 for a full explanation). In all cases, the values obtained are well below *H* = 0.34, which is an intuitive reference point corresponding to the Hellinger distance between two Gaussian distributions of identical variance, *σ*^2^, and difference in mean given by, Δ*µ* = *σ*.

**Figure 5:**
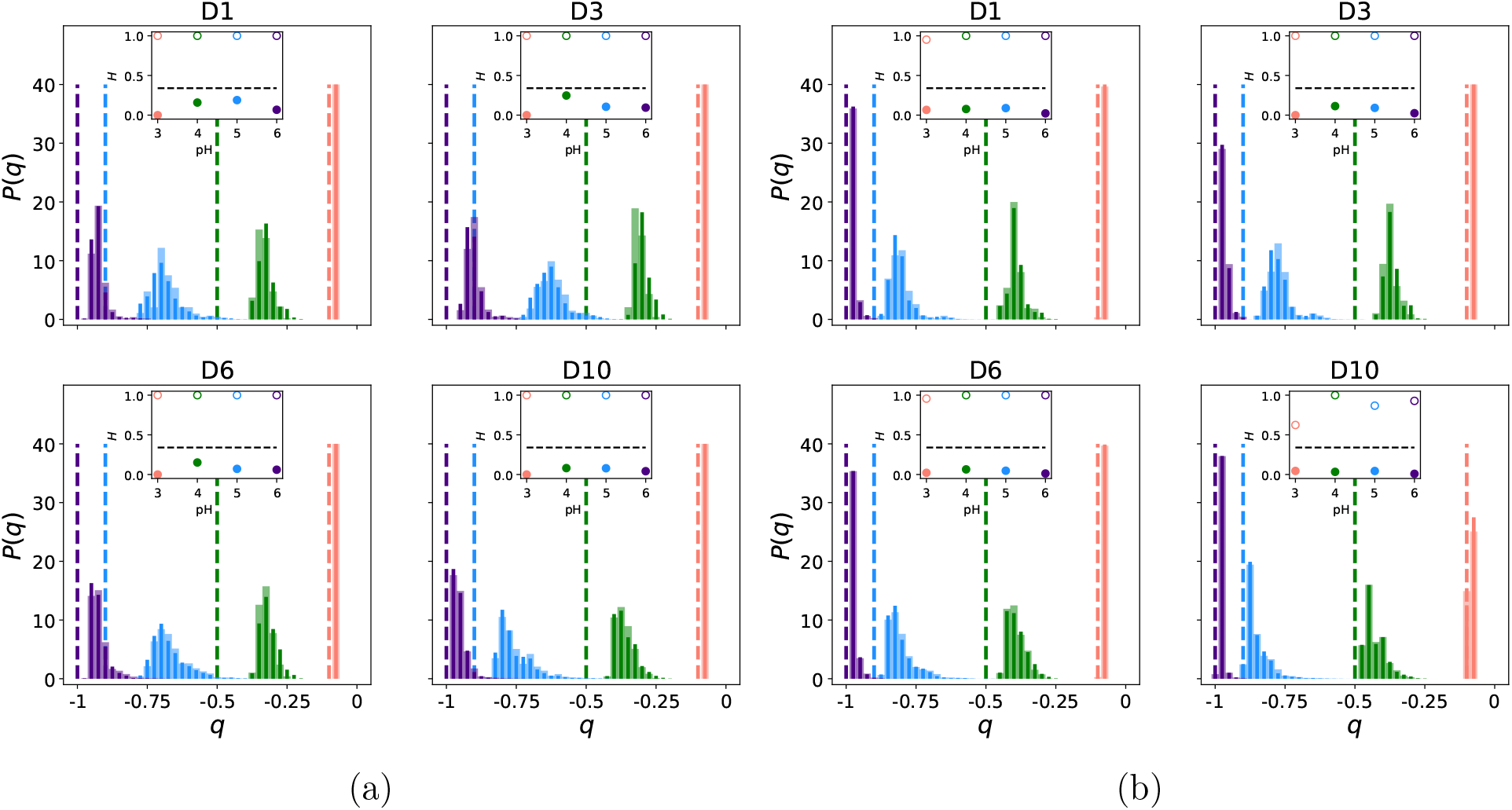
Charge distribution on each titratable group. In each figure, colors indicate the pH value: light-red is pH = 3, pH = 4 is in green, blue refers to pH = 5, and purple indicates pH = 6. Each figure is broken into 4 panels referring to the four titratable groups, D1, D3, D6, and D10. Wide, pale bars refer to the exact distribution; thinner, darker lines pertain to the MF approximation; dashed-bars refer to FC results. The insets show the Hellinger distances (see Eq. 27) between the exact and the MF results (full symbols) and between the exact and FC distributions (empty dots). In the insets, the dashed lines are to guide intuition: they are the values of *H* obtained for two Gaussian distributions of identical variance *σ*, and different means *µ*_1_ − *µ*_2_ = *σ*. The two figures refer to different salt concentrations: (a) 1 mM, (b) 100 mM.

The comparison with the FC results is instructive: not only the breadth of the distribution, but also its location are significantly altered by electrostatic interactions. Even at the largest ionic strength considered, the comparison with the FC results showcases the different distributions on the 4 aspartic acids, which are distinguished by the structure of the chain around them.

## Discussion

We have developed a practical method, based on theoretical considerations, to examine pH effects not only on IDPs but also any molecule (for example polyelectrolyte) with many titratable groups. The theory, which utilizes a MF approximation for treating titratable groups and the Molecular Transfer Model for summing over conformations, is tested using a lattice model for which numerically exact computations can be performed. The results show that, except at very low salt concentrations, the MF and MTM combination produces very accurate results. We close with a few pertinent comments.

- Although it has been suspected that the MF approximation leads to systematic errors in the estimate of the titration of acidic and basic groups in folded globular proteins with a fixed native structure,^14,15^ the accuracy of the MF for IDPs had not been previously studied. Our premise is that the MF approximation would be not be severe in disordered polymers that undergoe large conformational fluctuations. Two arguments support this assertion. First, the lack of a fixed structure reduces the correlation between two titratable groups: for all conformations in which these two residues are close
- enough to affect each other’s ionization, there are many other structures explored by the polymer in which the groups are further apart, and are likely to undergo titration independently. Second, in contrast to globular proteins, which present fully desolvated anionic and cationic groups (for instance in the vicinity of the active sites), IDPs are expanded, and it is reasonable to expect that all the ionizable groups are at least partially solvated in the majority of the conformations. As a consequence, electrostatic interactions in the IDPs are likely to be weakened by the dielectric permittivity of water and by ionic screening. Ultimately, both of these arguments stem from the idea that the presence of a heterogeneous ensemble of conformations mitigates the effects on average properties, which may not be appropriate if only a subset of conformations with specific interactions, such as the folded states, is considered.
- In order to test our assertion, we constructed a IDP-like polymer on a lattice, whose conformations can be enumerated exhaustively, thus avoiding the sampling issues. We used a sequence with 40% of charged beads, which is high but not unrealistic for many IDPs. The small number of ionizable groups enables the exact evaluation of the model partition function, which can be used to assess the accuracy of the MF estimate of the transfer free energy. We considered observables that depend on either the structure (radius of gyration and contact map) or the ionization state (average charge and charge correlation) of the polymer. We found that for all of these observables the MF MTM-IDP method recovers the exact results accurately. The discrepancies do not deviate from the exact value by more than a few percentage points.
- The inconsistencies between the MF and exact results are larger for pH values corresponding to the swelling (collapse) or charging (uncharging) of the polymer – when the pH is much larger or much smaller than the pKa of the titrating group, a single protonation state is effectively dominant. In agreement with our conjectures, as the salt concentration is increased the agreement between the MF and exact results gets better. At very low salt concentration, the MF approximation of the transfer free
- energy breaks down, as shown in Fig. 2. This problem may be exacerbated if the charge density along the chain is higher than what we used here, although it is likely to depend on the precise sequence. Although we have not explored the dependence on charge density and distribution systematically, we expect, as in the case reported here, that increasing the salt concentration is likely to diminish these issues.
- An important finding of this work is that the precise sequence of the IDP, especially the location of the charged residues, affects many observables. Properties of the IDPs could change drastically depending on charge correlations along the sequence.^43,44^ This is explicitly demonstrated in a model that does not suffer from sampling issues, and for which numerically exact results may be calculated. This finding shows that conclusions based on the overall net charge or polyampholytic parameter (charge asymmetry, calculable from charge composition alone without regard to sequence location of IDP conformations) cannot correctly produce the phase behavior of IDPs or how they might change as a function of pH or denaturants.
- It is important to note that fixing the charges along the chain at their pH-dependent average value results in a poor recovery of the exact results at all salt concentrations, except at high values that far exceeds the physiological value. This is interesting because it shows that although the chain is expanded, and the ionized groups are well-solvated, interactions between charged groups are important. It was already noted that this simple model lacking structural effects on titration frequently gives results close to the experimental values, and shows that the most dramatic deviations occur when strongly interacting groups are present.^17^ Here, we showed that although there are no de-solvated pairs, the simple model fails in reproducing properties such as the slope (co-operativity) and the mid-point of the polymer swelling as a function of pH.

The theory and applications show that the MTM-IDP method with MF approximation provides an accurate framework to estimate the change in the structural ensemble of an IDP as a function of solution pH. The applications of the method for lattice models for which exact calculations can be performed, thus allowing us to assess the MF approximation, demonstrates the success of the theory. The MTM-IDP method is general that it can be used to investigate pH effects in a number of biological as well as synthetic systems.

## Acknowledgement

This work was supported by National Science Foundation (CHE 19-00093), National Institute of Health (GM-107703), the Welch Foundation (F-0019) administered through the Collie-Welch chair.

## References

(1) Paroutis, P., Touret, N., Grinstein, S. The pH of the Secretory Pathway: Measurement, Determinants, and Regulation. Physiology (Bethesda, Md.) 2004, 19, 207–215.

(2) Bhatnagar, V., Anjaiah, S., Puri, N., Darshanam, B., Ramaiah, A. pH of Melanosomes of B 16 Murine Melanoma Is Acidic: Its Physiological Importance in the Regulation of Melanin Biosynthesis. Archives of biochemistry and biophysics 1993, 307, 183–192.

(3) Santo-Domingo, J., Demaurex, N. The renaissance of mitochondrial pH. The Journal of General Physiology 2012, 139, 415–423.

(4) Munder, M. C., Midtvedt, D., Franzmann, T., Nüske, E., Otto, O., Herbig, M., Ulbricht, E., Müller, P., Taubenberger, A., Maharana, S., Malinovska, L., Richter, D., Guck, J., Zaburdaev, V., Alberti, S. A pH-driven transition of the cytoplasm from a fluid-to a solid-like state promotes entry into dormancy. eLife 2016, 5.

(5) Love, C., Steinkühler, J., Gonzales, D. T., Yandrapalli, N., Robinson, T., Dimova, R., Tang, T.-Y. D. Reversible pH-Responsive Coacervate Formation in Lipid Vesicles Activates Dormant Enzymatic Reactions. Angewandte Chemie (International ed.) 2020, 59, 5950–5957.

(6) Last, M. G. F., Deshpande, S., Dekker, C. pH-Controlled Coacervate–Membrane Interactions within Liposomes. ACS nano 2020, 14, 4487–4498.

(7) Franzmann, T. M., Jahnel, M., Pozniakovsky, A., Mahamid, J., Holehouse, A. S., Nüske, E., Richter, D., Baumeister, W., Grill, S. W., Pappu, R. V., Hyman, A. A., Alberti, S. Phase separation of a yeast prion protein promotes cellular fitness. Science (American Association for the Advancement of Science) 2018, 359, eaao5654.#x2013;.

(8) Uversky, V. N., Li, J., Fink, A. L. Evidence for a partially folded intermediate in α-synuclein fibril formation. Journal of Biological Chemistry 2001, 276, 10737–10744.

(9) McGlinchey, R. P., Jiang, Z., Lee, J. C. Molecular Origin of pH-Dependent Fibril Formation of a Functional Amyloid. Chembiochem : a European journal of chemical biology 2014, 15, 1569–1572.

(10) Dogra, P., Joshi, A., Majumdar, A., Mukhopadhyay, S. Intermolecular Charge-Transfer Modulates Liquid–Liquid Phase Separation and Liquid-to-Solid Maturation of an Intrinsically Disordered pH-Responsive Domain. Journal of the American Chemical Society 2019, 141, 20380–20389.

(11) Malay, A. D., Suzuki, T., Katashima, T., Kono, N., Arakawa, K., Numata, K. Spider silk self-assembly via modular liquid-liquid phase separation and nanofibrillation. Science Advances 2020, 6.

(12) Tanford, C., Kirkwood, J. G. Theory of protein titration curves. I. General equations for impenetrable spheres. Journal of the American Chemical Society 1957, 79, 5333–5339.

(13) Baul, U., Chakraborty, D., Mugnai, M. L., Straub, J. E., Thirumalai, D. Sequence effects on size, shape, and structural heterogeneity in Intrinsically Disordered Proteins. The Journal of Physical Chemistry B 2019, 123, 3462–3474.

(14) Bashford, D., Karplus, M. Multiple-site titration curves of proteins: an analysis of exact and approximate methods for their calculation. The Journal of Physical Chemistry 1991, 95, 9556–9561.

(15) Gilson, M. K. Multiple-site titration and molecular modeling: Two rapid methods for computing energies and forces for ionizable groups in proteins. Proteins: Structure, Function, and Bioinformatics 1993, 15, 266–282.

(16) Mertz, J. E., Pettitt, B. M. Molecular Dynamics At a Constant pH. The international journal of supercomputer applications and high performance computing 1994, 8, 47–53.

(17) Antosiewicz, J., McCammon, J. A., Gilson, M. K. Prediction of Ph-dependent Properties of Proteins. Journal of Molecular Biology 1994, 238, 415–436.

(18) Baptista, A. M., Martel, P. J., Petersen, S. B. Simulation of protein conformational freedom as a function of pH: constant-pH molecular dynamics using implicit titration. Proteins: Structure, Function, and Bioinformatics 1997, 27, 523–544.

(19) Baptista, A. M., Teixeira, V. H., Soares, C. M. Constant-pH molecular dynamics using stochastic titration. The Journal of chemical physics 2002, 117, 4184–4200.

(20) Bürgi, R., Kollman, P. A., van Gunsteren, W. F. Simulating proteins at constant pH: An approach combining molecular dynamics and Monte Carlo simulation: Simulating Proteins at Constant pH. Proteins, structure, function, and bioinformatics 2002, 47, 469–480.

(21) Lee, M. S., Salsbury, F. R., Brooks, C. L. Constant-pH molecular dynamics using continuous titration coordinates. Proteins: Structure, Function, and Bioinformatics 2004, 56, 738–752.

(22) Mongan, J., Case, D. A., McCammon, J. A. Constant pH molecular dynamics in generalized Born implicit solvent. Journal of Computational Chemistry 2004, 25, 2038–2048.

(23) Stern, H. A. Molecular simulation with variable protonation states at constant p H. Journal of Chemical Physics 2007, 126, 164112–164112–7.

(24) Itoh, S. G., Damjanović, A., Brooks, B. R. pH replica-exchange method based on discrete protonation states. Proteins: Structure, Function, and Bioinformatics 2011, 79, 3420–3436.

(25) Swails, J. M., York, D. M., Roitberg, A. E. Constant pH Replica Exchange Molecular Dynamics in Explicit Solvent Using Discrete Protonation States: Implementation, Testing, and Validation. Journal of Chemical Theory and Computation 2014, 10, 1341– 1352.

(26) Chen, Y., Roux, B. Constant-pH Hybrid Nonequilibrium Molecular Dynamics–Monte Carlo Simulation Method. Journal of chemical theory and computation 2015, 11, 3919– 3931.

(27) Radak, B. K., Chipot, C., Suh, D., Jo, S., Jiang, W., Phillips, J. C., Schulten, K., Roux, B. Constant-pH Molecular Dynamics Simulations for Large Biomolecular Systems. Journal of chemical Theory and Computation 2017, 13, 5933–5944.

(28) O’Brien, E. P., Ziv, G., Haran, G., Brooks, B. R., Thirumalai, D. Effects of denaturants and osmolytes on proteins are accurately predicted by the molecular transfer model. Proceedings of the National Academy of Sciences 2008, 105, 13403–13408.

(29) Liu, Z., Reddy, G., Thirumalai, D. Theory of the Molecular Transfer Model for Proteins with Applications to the Folding of the src-SH3 Domain. The journal of physical chemistry. B 2012, 116, 6707–6716.

(30) O’Brien, E. P., Brooks, B. R., Thirumalai, D. Effects of pH on proteins: predictions for ensemble and single-molecule pulling experiments. Journal of the American Chemical Society 2012, 134, 979–987.

(31) Enciso, M., Schtte, C., Delle Site, L. A pH-dependent coarse-grained model for peptides. Soft matter 2013, 9, 6118–6127.

(32) Barroso da Silva, F.L., Sterpone, F., Derreumaux, P. OPEP6: A New Constant-pH Molecular Dynamics Simulation Scheme with OPEP Coarse-Grained Force Field. Journal of Chemical Theory and Computation 2019, 15, 3875–3888.

(33) Grünewald, F., Souza, P. C. T., Abdizadeh, H., Barnoud, J., de Vries, A. H., Marrink, S. J. Titratable Martini model for constant pH simulations. The Journal of chemical physics 2020, 153, 24118.

(34) Vo, P., Forsman, J., Woodward, C. E. A semi-GCMC simulation study of electrolytic capacitors with adsorbed titrating peptides. The Journal of chemical physics 2020, 153, 174703–174703.

(35) Tanford, C., Roxby, R. Interpretation of protein titration curves. Application to lysozyme. Biochemistry 1972, 11, 2192–2198.

(36) Hill, T. L. An Introdcution to Statistical Thermodynamics; Dover Publications, Inc.: New York, 1986.

(37) Nozaki, Y., Tanford, C. Examination of titration behavior. 1967.

(38) Klimov, D., Thirumalai, D. Mechanisms and kinetics of beta-hairpin formation. Proc. Natl. Acad. Sci. 2000, 97, 2544–2549.

(39) Wu, H., Wolynes, P. G., Papoian, G. A. AWSEM-IDP: A Coarse-Grained Force Field for Intrinsically Disordered Proteins. J. Phys. Chem. 2018, 122, 11115–11125.

(40) Latham, A. P., Zhang, B. Maximum Entropy Optimized Force Field for Intrinsically Disordered Proteins. J. Chem. Theor. Comp. 2020, 16, 773–781.

(41) Dill, K., Bromberg, S., Yue, K., Fiebig, K., Yee, D., Thomas, P., Chan, H. Principles of Protein-Folding-A Perspective From Simple Exact Models. Prot. Sci. 1995, 4, 561–602.

(42) Fisher, M. E., Sykes, M. F. Excluded-Volume Problem and the Ising Model of Ferromagnetism. Physical review 1959, 114, 45–58.

(43) Samanta, H. S., Chakraborty, D., Thirumalai, D. Charge fluctuation effects on the shape of flexible polyampholytes with applications to intrinsically disordered proteins. The Journal of chemical physics 2018, 149, 163323–163323.

(44) Firman, T., Ghosh, K. Sequence charge decoration dictates coil-globule transition in intrinsically disordered proteins. J. Chem. Phys. 2018, 148, 123305.

