## Supplementary Information for "Molecular Transfer Model for pH effects on Intrinsically Disordered Proteins: Theory and Applications"

A crucial component in the MTM-IDP method is the determination of the transfer free energy, $\Delta G_{\rm tr}(c, \alpha^\star, {\rm pH})$, given in Eq. 15 in the text. We show that the fairly general expression (Eq. 15) reduces to equations derived previously[S1,S2] with appropriate choices of the reference states. With some effort we also show that Eq. 15 could be used to derive the equation used for computing the pH dependence of the stability changes in globular proteins.[S3]

#### Comparison with Previous Studies

**Comparison with Gilson:[S1]** We begin with Eq. 15 in the main text, and set $x_i(\alpha^\star) = 0$ for all the titratable groups, which corresponds to the fully de-ionized reference state used by Gilson. Next, we consider the exponent in curly brackets in Eq. 15, which is given by a product of two terms:

$$e^{-\beta\sum_i x_i(\alpha)[b_i+\frac{1}{2}\sum_j x_j(\alpha)V_{ij}(c)]} = e^{-\beta\sum_i b_i x_i(\alpha)} e^{-\beta\sum_i\sum_{j>i} V_{ij}x_i(\alpha)x_j(\alpha)}. \tag{S1}$$

The second term on the r.h.s. is equivalent to $G_{\text{elec}}(\alpha)$ (note that Gilson uses $\alpha$ as a subscript, but this should not create any confusion) in Eq. 14 of Gilson's work.[S1] The only difference is the presence of a term $G_i$ in the exponent in Eq. 14,[S1] which we have absorbed into the definition of the equilibrium dissociation constant. We then "unpack" $b_i$ in Eq. 11, and write:

$$e^{-\beta\sum_i x_i(\alpha)b_i} = e^{\sum_i z_i x_i(\alpha)\ln(10)(\text{pKa}_i-\text{pH})} = \prod_i \left(\frac{[\text{H}^+]}{K_{\text{a},i}}\right)^{z_i x_i(\alpha)}, \tag{S2}$$

where we used the definitions of pH and pKa of the $i$-th group. Gilson's Eq. 15 introduces the definition $A_i = K_{\text{a},i}^{-z_i}[\text{H}^+]^{z_i}$; we plug this and $G_{\text{elec}}(\alpha)$ in Eq. 15 and obtain,

$$\Delta G_{\text{tr}} = -k_B T \ln\Big\{\sum_\alpha e^{-\beta G_{\text{elec}}(\alpha)} \prod_i A_i^{x_i(\alpha)}\Big\}, \tag{S3}$$

which is the same as Eq. 16 in Gilson's work.[S1]

**Comparison with Bashford-Karplus:[S2]** Bashford and Karplus (BK) used protonation states, $p_i(\alpha)$, instead of ionization states, $x_i(\alpha)$ (they used $x$ for protonation state, we introduce $p$ to avoid any confusion, the relationship between the two is given in Eq. 1). Furthermore, BK considered a reference state in which all titratable groups are de-protonated, that is

$$x_i(\alpha^\star) = z_i q_i^{\text{unp}}. \tag{S4}$$

For convenience, we rewrite Eq. 15 as,

$$\Delta G_{\text{tr}}(c,\alpha^\star,\text{pH}) = -k_B T \ln\Big\{\sum_\alpha e^{-\beta\Delta G_{\text{tr}}(c,\alpha,\alpha^\star,\text{pH})}\Big\}, \tag{S5}$$

and we replace in $\Delta G_{\text{tr}}(c,\alpha,\alpha^\star,\text{pH})$ the ionization state with the protonation state, and we introduce the appropriate reference state. These transformations lead to,

$$
\begin{aligned}
\Delta G_{\text{tr}}(c,\alpha,\alpha^\star,\text{pH}) &= \sum_{i=1}^{N} b_i[x_i(\alpha)-x_i(\alpha^\star)] + \frac{1}{2}\sum_{i=1}^{N}\sum_{j=1}^{N} V_{ij}[x_i(\alpha)x_j(\alpha)-x_i(\alpha^\star)x_j(\alpha^\star)] = \\
&= \sum_{i=1}^{N} b_i[x_i(\alpha)-z_i q_i^{\text{unp}}] + \frac{1}{2}\sum_{i=1}^{N}\sum_{j=1}^{N} V_{ij}[x_i(\alpha)x_j(\alpha)-z_i z_j q_i^{\text{unp}} q_j^{\text{unp}}] = \\
&= \sum_{i=1}^{N} b_i z_i p_i(\alpha) + \frac{1}{2}\sum_{i=1}^{N}\sum_{j=1}^{N} V_{ij} z_i z_j [p_i(\alpha)p_j(\alpha)+p_i(\alpha)q_j^{\text{unp}}+p_j(\alpha)q_i^{\text{unp}}] = \\
&= \sum_{i=1}^{N}(b_i z_i + \sum_{j=1}^{N} W_{ij} q_j^{\text{unp}}) p_i(\alpha) + \frac{1}{2}\sum_{i=1}^{N}\sum_{j=1}^{N} W_{ij} p_i(\alpha) p_j(\alpha).
\end{aligned}
\tag{S6}
$$

where we wrote $V_{ij}z_iz_j = W_{ij}$, where $W_{ij}$ is now an electrostatic interaction between two unit charges, as in Eq. 1a of BK.[S2] In order to complete the comparison, we note that,

$$
b_i z_i + \sum_{j=1}^{N} W_{ij} q_j^{\text{unp}} = -k_B T(\ln 10)(\text{pKa}_i - \text{pH}) + \sum_{j=1}^{N} W_{ij} q_j^{\text{unp}}, \tag{S7}
$$

which is exactly the same as the $b_i$ defined in Eq. 8 of BK.[S2] Therefore,

$$
b_i^{\text{BK}} = -k_B T(\ln 10)(\text{pKa}_i - \text{pH}) + \sum_{j=1}^{N} W_{ij} q_j^{\text{unp}}. \tag{S8}
$$

Using these transformations, we obtain from Eq. 15,

$$
\Delta G_{\text{tr}} = -k_B T \ln \Big\{ \sum_{\alpha} e^{-\beta \sum_{i=1}^{N}[b_i^{\text{BK}} p_i(\alpha) + \frac{1}{2}\sum_{j=1}^{N} W_{ij} p_i(\alpha) p_j(\alpha)]} \Big\}. \tag{S9}
$$

From this expression, we can obtain $\Delta G_{\text{tr}}$ using the same density functional minimization used by BK and obtain,

$$
\Delta G_{\text{tr}} = \min_{\rho} \sum_{\alpha} \Big\{ \sum_{i=1}^{N} b_i^{\text{BK}} p_i(\alpha) + \frac{1}{2}\sum_{j=1}^{N} W_{ij} p_i(\alpha) p_j(\alpha) + k_B T \ln \rho[\vec{p}(\alpha)] \Big\} \rho[\vec{p}(\alpha)], \tag{S10}
$$

which is identical to Eq. 9 of BK.[S2]

**Comparison with O'Brien, Brooks, and Thirumalai:**[S3] We start again from Eq. 15, and we attempt to recover Eq. 7 of O'Brien, Brooks, and Thirumalai (OBT).[S3] There are two key points here. First, OBT compute the transfer free energy for going from a solution at a given pH, to a second solution at a different pH. This differs from Eq. 15, which is the free energy for equilibrating a fixed-charge conformation at a given pH. The idea is to apply a re-weighting of the conformations sampled at fixed protonation state to model an ensemble at $\mathrm{pH}_1$, and then to define a second transformation that allows to change from $\mathrm{pH}_1$ to $\mathrm{pH}_2$, which will be related to Eq. 7 of OBT.[S3] Using Eq. 8 we define,

$$\begin{aligned}
&\Delta\Delta G_{\mathrm{tr}}(c, \mathrm{pH}_1 \to \mathrm{pH}_2) = \\
&\quad \Delta G_{\mathrm{tr}}(c, \alpha^\star, \mathrm{pH}_2) + E_1(\alpha^\star) - \mu_{\mathrm{H},2}\nu(\alpha^\star) - \Delta G_{\mathrm{tr}}(c, \alpha^\star, \mathrm{pH}_1) - E_1(\alpha^\star) + \mu_{\mathrm{H},1}\nu(\alpha^\star) = \\
&\quad = \Delta G_{\mathrm{tr}}(c, \alpha^\star, \mathrm{pH}_2) - \Delta G_{\mathrm{tr}}(c, \alpha^\star, \mathrm{pH}_1) - (\mu_{\mathrm{H},2} - \mu_{\mathrm{H},1})\nu(\alpha^\star) = \\
&\quad = \Delta G_{\mathrm{tr}}(c, \alpha^\star, \mathrm{pH}_2) - \Delta G_{\mathrm{tr}}(c, \alpha^\star, \mathrm{pH}_1) + \nu(\alpha^\star)\ln(10)(\mathrm{pH}_2 - \mathrm{pH}_1).
\end{aligned} \tag{S11}$$

Second, in the OBT model the proton dissociation constant (thus, the pKa) of each titratable group depends on the configuration, but not on the ionization state of the other cations and anions. This means that, compared to the definitions adopted here, we must modify $\Delta G_{\mathrm{tr}}(c, \alpha^\star, \mathrm{pH})$. First, we "unpack" Eq. 15 by writing explicitly $b_i$ (see Eq. 11),

$$\begin{aligned}
&\Delta G_{\mathrm{tr}}(c, \alpha^\star, \mathrm{pH}) = -k_B T \ln\Big\{\sum_\alpha e^{-\sum_i \ln(10) z_i x_i(\alpha)\mathrm{pH}} e^{\sum_i z_i x_i(\alpha)[\ln(10)\mathrm{pKa}_i - \sum_{j>i}\beta W_{ij}(c) z_j x_j(\alpha)]}\Big\} + \\
&- \sum_i k_B T \ln(10) z_i x_i(\alpha^\star)\mathrm{pH} + \sum_i k_B T z_i x_i(\alpha^\star)[\ln(10)\mathrm{pKa}_i - \sum_{j>i} W_{ij}(c) z_j x_j(\alpha^\star)],
\end{aligned} \tag{S12}$$

where as before $V_{ij} = z_i z_j W_{ij}$, with $W_{ij}$ being an interaction between the unit charges. Here, we introduce the feature that the pKa of a group depends only on the conformation and not on the protonation states. We do so, by replacing, $\ln(10)\mathrm{pKa}_i - \frac{1}{2}\sum_j W_{ij}(c) z_j x_j(\alpha)$, with, $\ln(10)\mathrm{pKa}_i(c)$, where $\mathrm{pKa}_i(c)$ depends on the conformation but not on which group is ionized.

This qualitatively changes the model, and leads to a new expression for $\Delta G_{\text{tr}}(c, \alpha^\star, \text{pH})$,

$$\begin{aligned}\Delta G_{\text{tr}}(c, \alpha^\star, \text{pH}) = -k_B T \ln\Big\{\sum_\alpha e^{\sum_i \ln(10) z_i x_i(\alpha)[\text{pKa}_i(c) - \text{pH}]}\Big\} + \\ + k_B T \ln(10) \sum_i z_i x_i(\alpha^\star)[\text{pKa}_i(c) - \text{pH}].\end{aligned} \tag{S13}$$

Using the properties of the exponent we obtain,

$$\begin{aligned}\Delta G_{\text{tr}}(c, \alpha^\star, \text{pH}) = -k_B T \ln\Big\{\sum_\alpha \prod_i e^{\ln(10) z_i x_i(\alpha)[\text{pKa}_i(c) - \text{pH}]}\Big\} + \\ + k_B T \ln(10) \sum_i z_i x_i(\alpha^\star)[\text{pKa}_i(c) - \text{pH}].\end{aligned} \tag{S14}$$

Next, we make the following observation: given a generic function $f_i(x_i)$ that depends on the protonation state of the $i$-th group alone, we can show that,

$$\sum_{\alpha=1}^{2^{N_T}} \prod_{i=1}^{N_T} f_i[x_i(\alpha)] = \prod_{i=1}^{N_T} \sum_{x=0}^{1} f_i(x). \tag{S15}$$

To prove this expression, we begin with $N_T = 2$, where the l.h.s gives,

$$\begin{aligned}\sum_{\alpha=1}^{4} f_1[x_1(\alpha)] f_2[x_2(\alpha)] = f_1(0)f_2(0) + f_1(1)f_2(0) + f_1(0)f_2(1) + f_1(1)f_2(1) = \\ = [f_1(0) + f_1(1)][f_2(0) + f_2(1)],\end{aligned} \tag{S16}$$

which is clearly the same as the right hand side of Eq. S15. Next, assuming that the identity in Eq. S15 holds for an arbitrary $N_T$, we show that it holds for $N_T + 1$ as well.

$$\begin{aligned}\sum_{\alpha=1}^{2^{N_T+1}} \prod_{i=1}^{N_T+1} f_i[x_i(\alpha)] = \sum_{\alpha=1}^{2^{N_T+1}} f_{N_T+1}[x_{N_T+1}(\alpha)] \prod_{i=1}^{N_T} f_i[x_i(\alpha)] = \\ = f_{N_T+1}(0) \sum_{\alpha=1}^{2^{N_T}} \prod_{i=1}^{N_T} f_i[x_i(\alpha)] + f_{N_T+1}(1) \sum_{\alpha=1}^{2^{N_T}} \prod_{i=1}^{N_T} f_i[x_i(\alpha)] = \\ = [f_{N_T+1}(0) + f_{N_T+1}(1)] \sum_{\alpha=1}^{2^{N_T}} \prod_{i=1}^{N_T} f_i[x_i(\alpha)] = \prod_{i=1}^{2^{N_T+1}} \sum_{x=0}^{1} f_i(x),\end{aligned} \tag{S17}$$

where with the last equality we have used that Eq. S15 holds for $N_T$. This proves by induction Eq. S15, and using this equality and the properties of the logarithm we have from Eq. S14,

$$\begin{aligned}\Delta G_{\mathrm{tr}}(c, \alpha^\star, \mathrm{pH}) = -k_B T \sum_i \ln\Big\{\sum_{x=0}^{1} e^{\ln(10) z_i x [\mathrm{pKa}_i(c) - \mathrm{pH}]}\Big\} + \\ + k_B T \ln(10) \sum_i z_i x_i(\alpha^\star)[\mathrm{pKa}_i(c) - \mathrm{pH}].\end{aligned} \tag{S18}$$

Using again the properties of the logarithm and performing the summation in the logarithm we obtain,

$$\begin{aligned}\Delta G_{\mathrm{tr}}(c, \alpha^\star, \mathrm{pH}) = -k_B T \sum_i \ln\Big\{1 + 10^{z_i[\mathrm{pKa}_i(c) - \mathrm{pH}]}\Big\} + \\ + k_B T \ln(10) \sum_i z_i x_i(\alpha^\star)[\mathrm{pKa}_i(c) - \mathrm{pH}].\end{aligned} \tag{S19}$$

Next, Eq. S11 becomes,

$$\begin{aligned}\Delta\Delta G_{\mathrm{tr}}(c, \mathrm{pH}_1 \to \mathrm{pH}_2) = -k_B T \sum_i \ln\Big\{\frac{1 + 10^{z_i[\mathrm{pKa}_i(c) - \mathrm{pH}_2]}}{1 + 10^{z_i[\mathrm{pKa}_i(c) - \mathrm{pH}_1]}}\Big\} + \\ - k_B T \ln(10) \sum_i x_i(\alpha^\star)(\mathrm{pH}_2 - \mathrm{pH}_1) + k_B T \ln(10) \nu(\alpha^\star)(\mathrm{pH}_2 - \mathrm{pH}_1).\end{aligned} \tag{S20}$$

For a cation ($z_i = 1$), we rewrite the argument of the logarithm as follows,

$$\ln\Big\{\frac{1 + 10^{[\mathrm{pKa}_i(c) - \mathrm{pH}_2]}}{1 + 10^{[\mathrm{pKa}_i(c) - \mathrm{pH}_1]}}\Big\} = -\ln(10)(\mathrm{pH}_2 - \mathrm{pH}_1) + \ln\Big\{\frac{10^{\mathrm{pH}_2} + 10^{\mathrm{pKa}_i(c)}}{10^{\mathrm{pH}_1} + 10^{\mathrm{pKa}_i(c)}}\Big\}. \tag{S21}$$

For an anionic group ($z_i = -1$), we get,

$$\ln\Big\{\frac{1 + 10^{[\mathrm{pH}_2 - \mathrm{pKa}_i(c)]}}{1 + 10^{[\mathrm{pH}_1 - \mathrm{pKa}_i(c)]}}\Big\} = \ln\Big\{\frac{10^{\mathrm{pH}_2} + 10^{\mathrm{pKa}_i(c)}}{10^{\mathrm{pH}_1} + 10^{\mathrm{pKa}_i(c)}}\Big\}. \tag{S22}$$

Plugging these expressions in Eq. S20 we obtain,

$$
\begin{aligned}
\Delta\Delta G_{\rm tr}(c,\mathrm{pH}_1\rightarrow \mathrm{pH}_2) &= -k_BT\sum_i \ln\left\{\frac{10^{\mathrm{pH}_2}+10^{\mathrm{pKa}_i(c)}}{10^{\mathrm{pH}_1}+10^{\mathrm{pKa}_i(c)}}\right\}+\\
&- k_BT\ln(10)\sum_i x_i(\alpha^\star)(\mathrm{pH}_2-\mathrm{pH}_1)+k_BT\ln(10)(\mathrm{pH}_2-\mathrm{pH}_1)N_C+\\
&+ k_BT\ln(10)\nu(\alpha^\star)(\mathrm{pH}_2-\mathrm{pH}_1),
\end{aligned}
\tag{S23}
$$

where $N_C$ is the number of cationic groups. Finally, the term outside the logarithm can be re-organized with the following considerations: $-\sum_i x_i(\alpha^\star)+N_C$ can be rewritten as, $\sum_{i\in\mathrm{Cation}}[1-x_i(\alpha^\star)]-\sum_{i\in\mathrm{Anion}}x_i(\alpha^\star)$, where the first (second) sum extends over cationic (anionic) groups. These two summations yield $N_T-\nu(\alpha^\star)$, that is the total number of titratable groups that are not protonated in the reference state – this is because the ionization state of a cationic group is 1 if protonated and zero otherwise, whereas for an anion $x_i(\alpha^\star)=1$ if the residue is de-protonated. This leads us to,

$$
\begin{aligned}
\Delta\Delta G_{\rm tr}(c,\mathrm{pH}_1\rightarrow \mathrm{pH}_2) &= -k_BT\sum_i \ln\left\{\frac{10^{\mathrm{pH}_2}+10^{\mathrm{pKa}_i(c)}}{10^{\mathrm{pH}_1}+10^{\mathrm{pKa}_i(c)}}\right\}+\\
&+ k_BT\ln(10)[N_T-\nu(\alpha^\star)](\mathrm{pH}_2-\mathrm{pH}_1)+k_BT\ln(10)\nu(\alpha^\star)(\mathrm{pH}_2-\mathrm{pH}_1)=\\
&- k_BT\sum_i \ln\left\{\frac{10^{\mathrm{pH}_2}+10^{\mathrm{pKa}_i(c)}}{10^{\mathrm{pH}_1}+10^{\mathrm{pKa}_i(c)}}\right\}+k_BT\ln(10)N_T(\mathrm{pH}_2-\mathrm{pH}_1)=\\
&- k_BT\sum_i \ln\left\{\frac{10^{\mathrm{pH}_2}+10^{\mathrm{pKa}_i(c)}}{10^{\mathrm{pH}_1}+10^{\mathrm{pKa}_i(c)}}\right\}-N_T(\mu_{\mathrm{H},2}-\mu_{\mathrm{H},1})
\end{aligned}
\tag{S24}
$$

The first term on the r.h.s. is exactly Eq. 7 of OBT,[S3] the second term is a constant dependent on the change in proton chemical potential and the total number of titratable groups. This constant does not affect the average of observables dependent on the conformation. Thus, we have recovered the MTM model adopted by OBT.

### Implementation of the Method and Model

The code was written in Python3 using functions from NumPy[S4,S5] libraries. Data was analyzed using MatPlotLib[S6] and Jupyter[S7] notebooks.

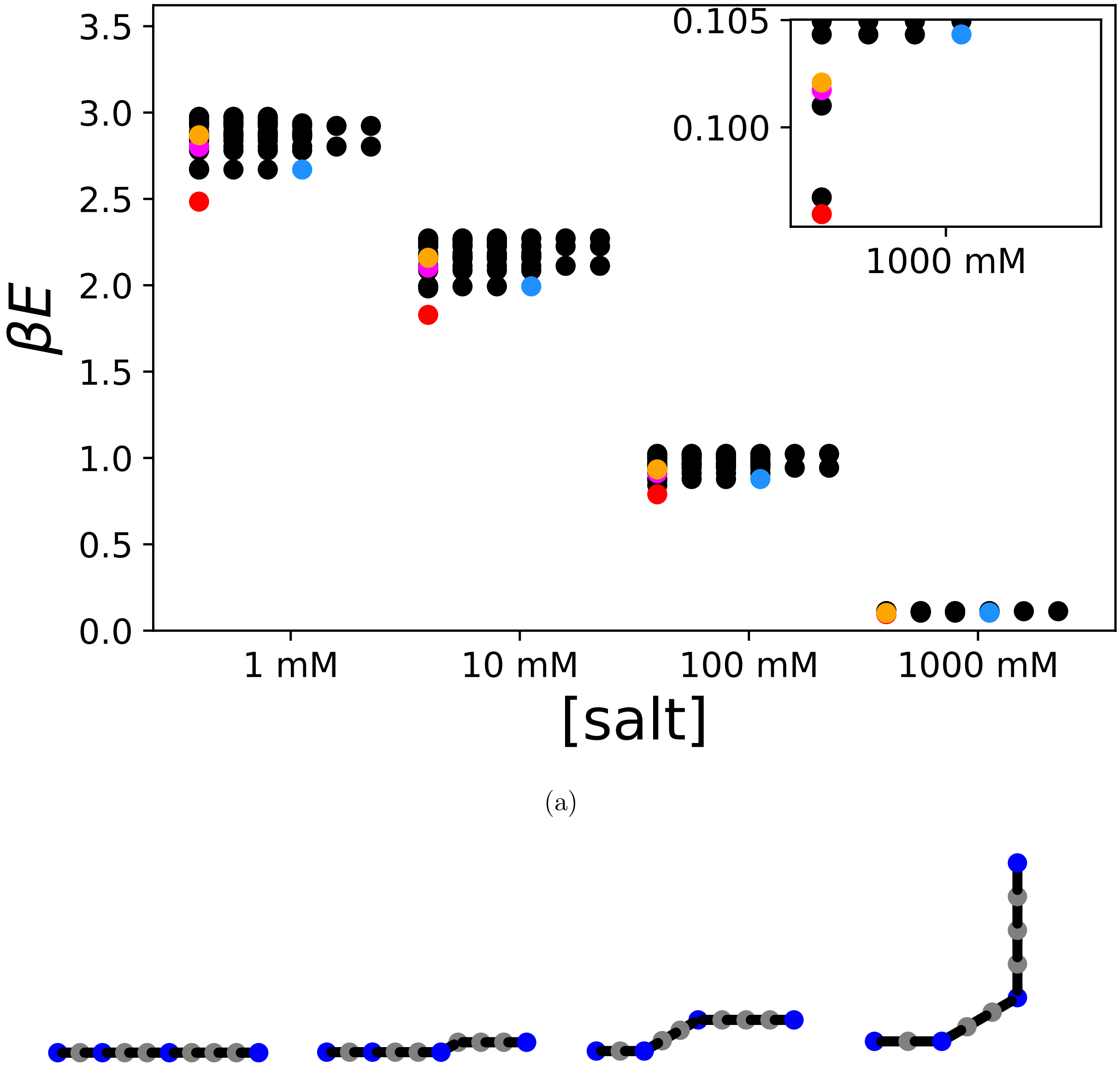


(a)

(b) red (c) blue (d) magenta (e) orange

Figure S1: Energy spectrum for the sequence DADAADAAAD. The aspartic acids are all charged, and the energy corresponds to the electrostatic interaction between anions. The 20 lowest energy levels are shown, with all the degenerate states for salt concentrations of 1 mM, 10 mM, 100 mM, 1000 mM. The inset highlights the results for ionic strength of 1000 mM. The red dot shows the ground state, the blue, magenta, and orange dots refer to two excited states. The structures for these four conformations are shown in panels (b-e) – the polymer is shown with gray dots for alanine residues and aspartic acids shown in blue.

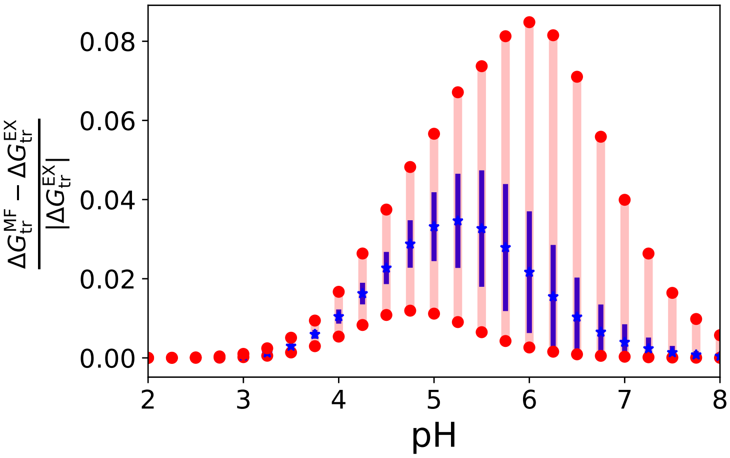


(a)

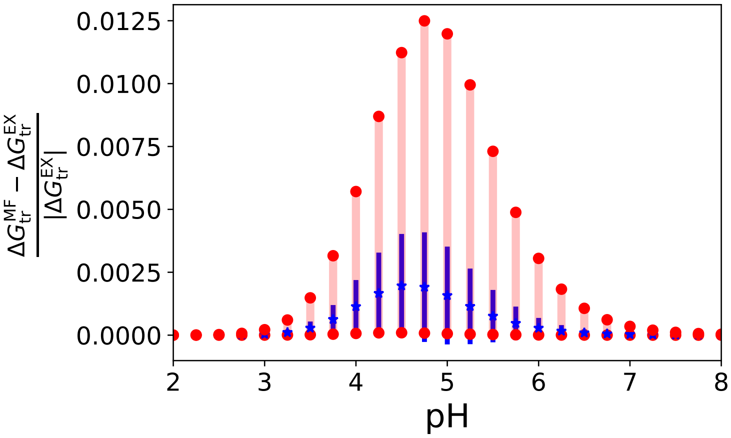


(b)

Figure S2: Comparison of the MF and exact calculations of the transfer free energy. Same as Fig. 2 in the main text, but for salt concentrations (a) 10 mM, (b) 1M.

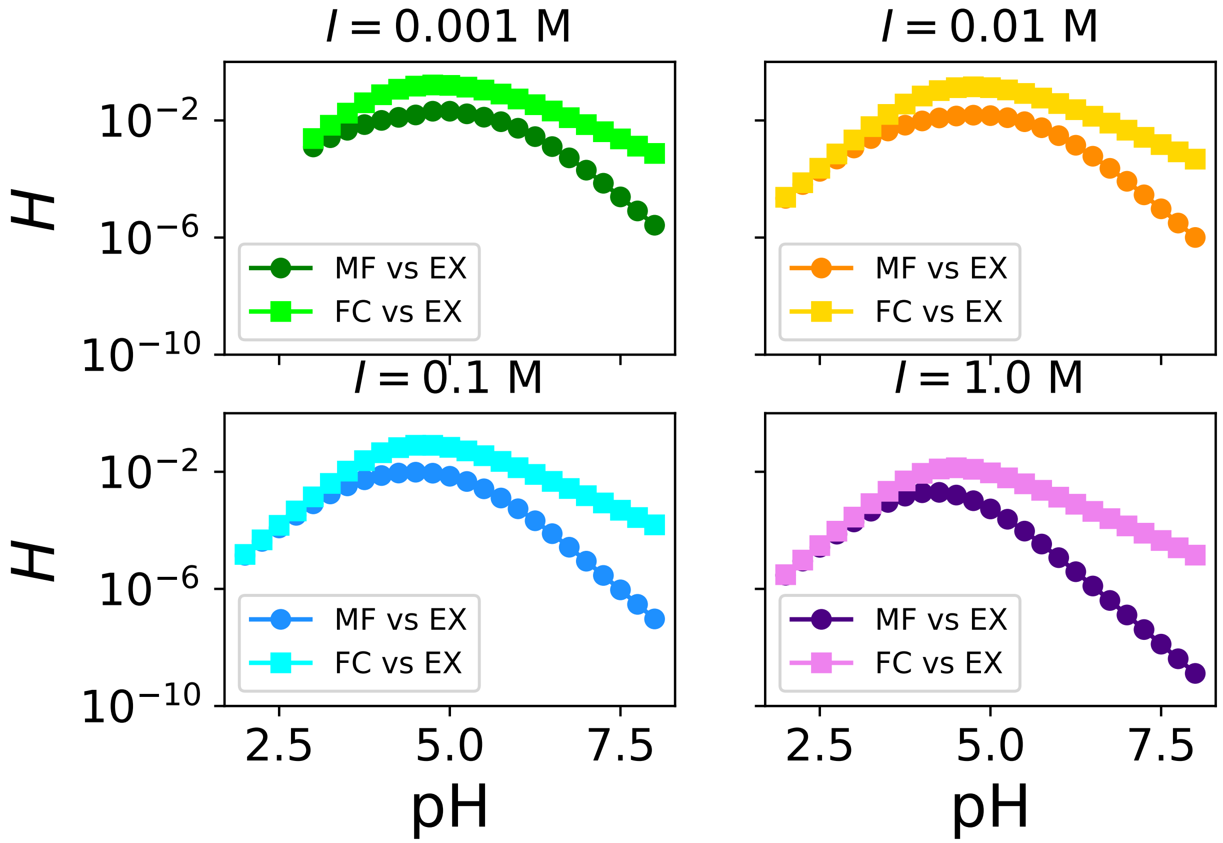


Figure S3: Comparison of the probability for each conformation. The four panels show the Hellinger distance (Eq. 27) for different ionic strengths (see numbers on top of each figure, from top left to bottom right the values are 1 mM, 10 mM, 100 mM, and 1 M). In each panel, the dark color (circles) refers to the comparison between the exact and MF probabilities, the light color (squares) report on the comparison between the exact and FC distributions.

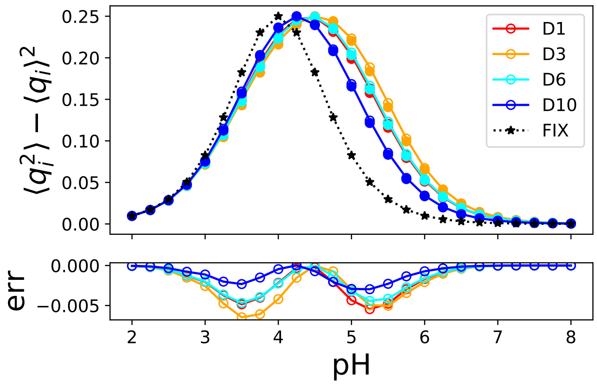


(a)

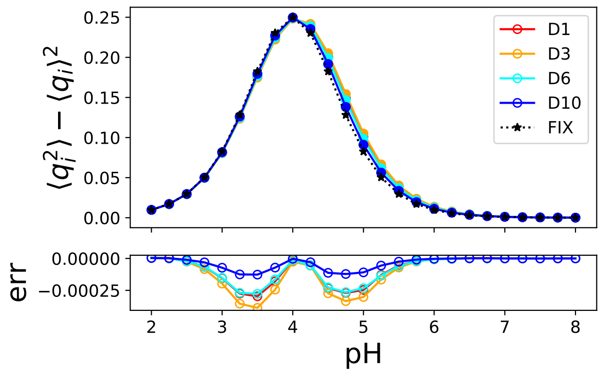


(b)

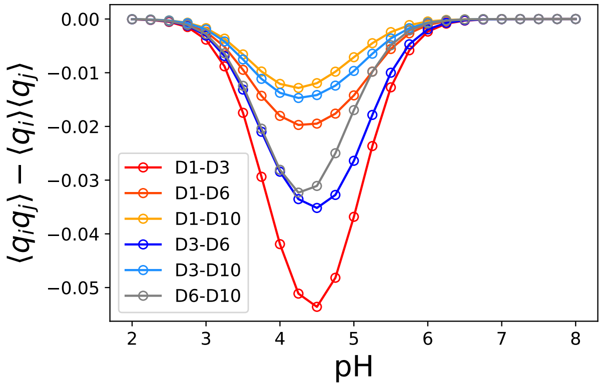


(c)

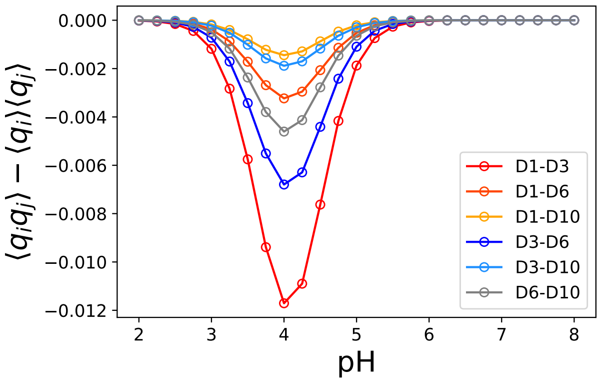


(d)

Figure S4: Comparison of the charge-charge correlation as a function of pH. Same as Fig. 4, but for salt concentrations (a) 10 mM, and (b) 1 M.

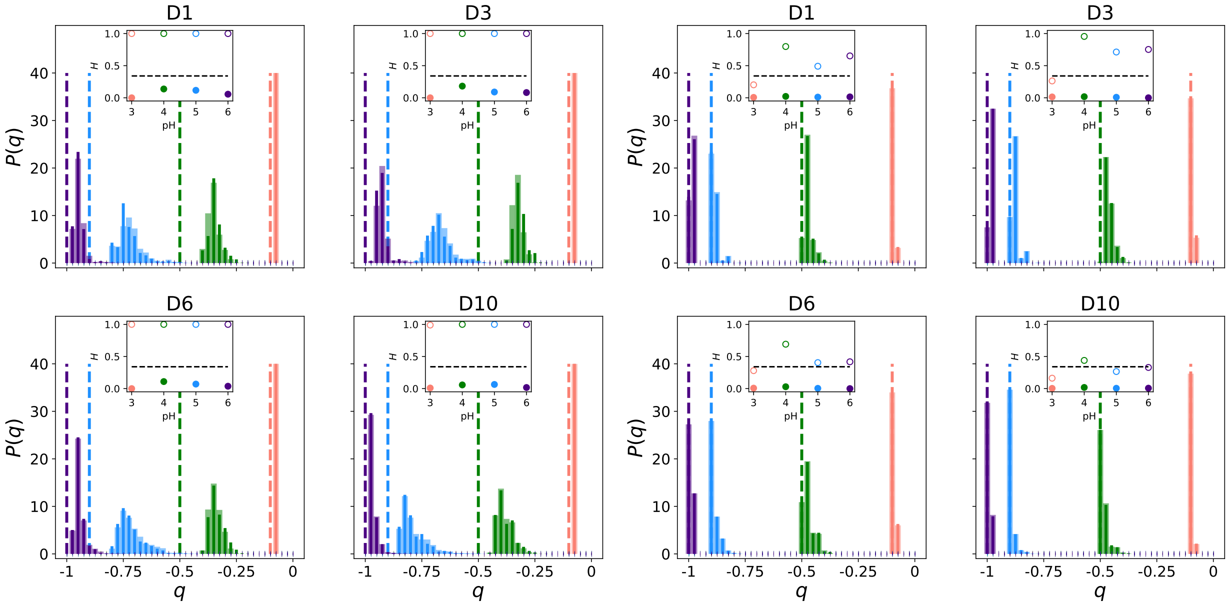


(a) (b)

Figure S5: Charge distribution on each titratable group. Same as Fig. 5, but for salt concentrations (a) 10 mM, (b) 1 M.

### References


(S1) Gilson, M. K. Multiple-site titration and molecular modeling: Two rapid methods for computing energies and forces for ionizable groups in proteins. *Proteins: Structure, Function, and Bioinformatics* **1993**, *15*, 266–282.

(S2) Bashford, D.; Karplus, M. Multiple-site titration curves of proteins: an analysis of exact and approximate methods for their calculation. *The Journal of Physical Chemistry* **1991**, *95*, 9556–9561.

(S3) O'Brien, E. P.; Brooks, B. R.; Thirumalai, D. Effects of pH on proteins: predictions for ensemble and single-molecule pulling experiments. *Journal of the American Chemical Society* **2012**, *134*, 979–987.

(S4) Oliphant, T. E. *A guide to NumPy*; Tregol Publishing, 2006.

(S5) van der Walt, S. J.; Colbert, S. C.; Gaël, V. The Numpy Array: A Structure for Efficient Numerical Computing. *Computing in Science & Engineering* **2011**, *13*, 22–30.

(S6) Hunter, J. D. Matplotlib: A 2D Graphics Environment. *Computing in Science & Engineering* **2007**, *9*, 90–95.

(S7) Kluyver, T.; Ragan-Kelley, B.; Pérez, F.; Granger, B.; Bussonnier, M.; Frederic, J.; Kelley, K.; Hamrick, J.; Grout, J.; Corlay, S.; Ivanov, P.; Avila, D.; Abdalla, S.; Willing, C.; Jupyter Development Team, Jupyter Notebooks-a publishing format for reproducible computational workflows. ELPUB. 2016; pp 87–90.
